# A novel role of circCPSF6 regulating antiviral innate immunity via miR-665 and PCBP2-IPS-1 axis

**DOI:** 10.1101/2025.11.03.686289

**Authors:** Aparna Meher, Riya Chaudhary, Himanshu Kumar

## Abstract

**Objective:** Circular RNAs are emerging as critical regulators of biological processes, yet their contribution to innate immune response during virus infection remains insufficiently defined. This study investigated the role of circCPSF6 in modulating host antiviral responses.

**Method:** CircCPSF6 expression was validated in human and murine cells and tissue by RT-PCR and digital PCR. Its effect on virus replication was analysed by RT-PCR in human and murine primary cells and cytokine production at protein level was measured by ELISA. Luciferase assay, RNA immunoprecipitation, RNA pulldown assay, immunoblotting was performed to explore the circRNA-miRNA regulatory mechanism. CircCPSF6 interacting protein were identified by RNA pulldown followed by mass spectrometry and regulatory mechanism was analysed by immunoblotting and confocal microscopy.

**Result:** CircCPSF6 identified as highly conserved circRNA, suppressed during virus infection, *in vitro* and *in vivo*. Functional analyses revealed antiviral role of circCPSF6. Mechanistically, circCPSF6 has dual cross-regulatory role by sequestering proviral miR-665 to relieve expression of key antiviral genes (MyD88, STAT2, IKKε) and interact with RNA binding protein PCBP2 to modulate IPS-1 degradation.

**Conclusion:** CircCPSF6 exert antiviral role and regulate host innate immune signaling by direct RNA-RNA and RNA-protein interaction, highlights circRNA mediated network as a potential therapeutic target in virus infection.

**Highlights:** CircCPSF6 is evolutionarily conserved and suppressed during virus infection.

CircCPSF6 act as antiviral factor by promoting inflammatory cytokine production.

CircCPSF6 sponges proviral miR-665 and alleviate MyD88, STAT2, IKKε expression.

CircCPSF6 bind to PCBP2, limit IPS-1 degradation and sustain antiviral signaling.

**GRAPHICAL ABSTRACT:** **Mechanism of host innate immune regulation by circCPSF6 during virus infection.** In cell cytoplasm circCPSF6 act as ceRNA for miR-665 and RNA binding protein PCBP2. Virus infection leads to reduced circCPSF6 and increased miR-665 expression, which inhibits expression of antiviral genes, MyD88, STAT2, IKKε. Additionally, reduced circCPSF6 leads to elevated free PCBP2 protein, hence, promote PCBP2 mediated IPS-1 degradation. Overall, these effects contribute to reduced innate immune response and increased viral replication.

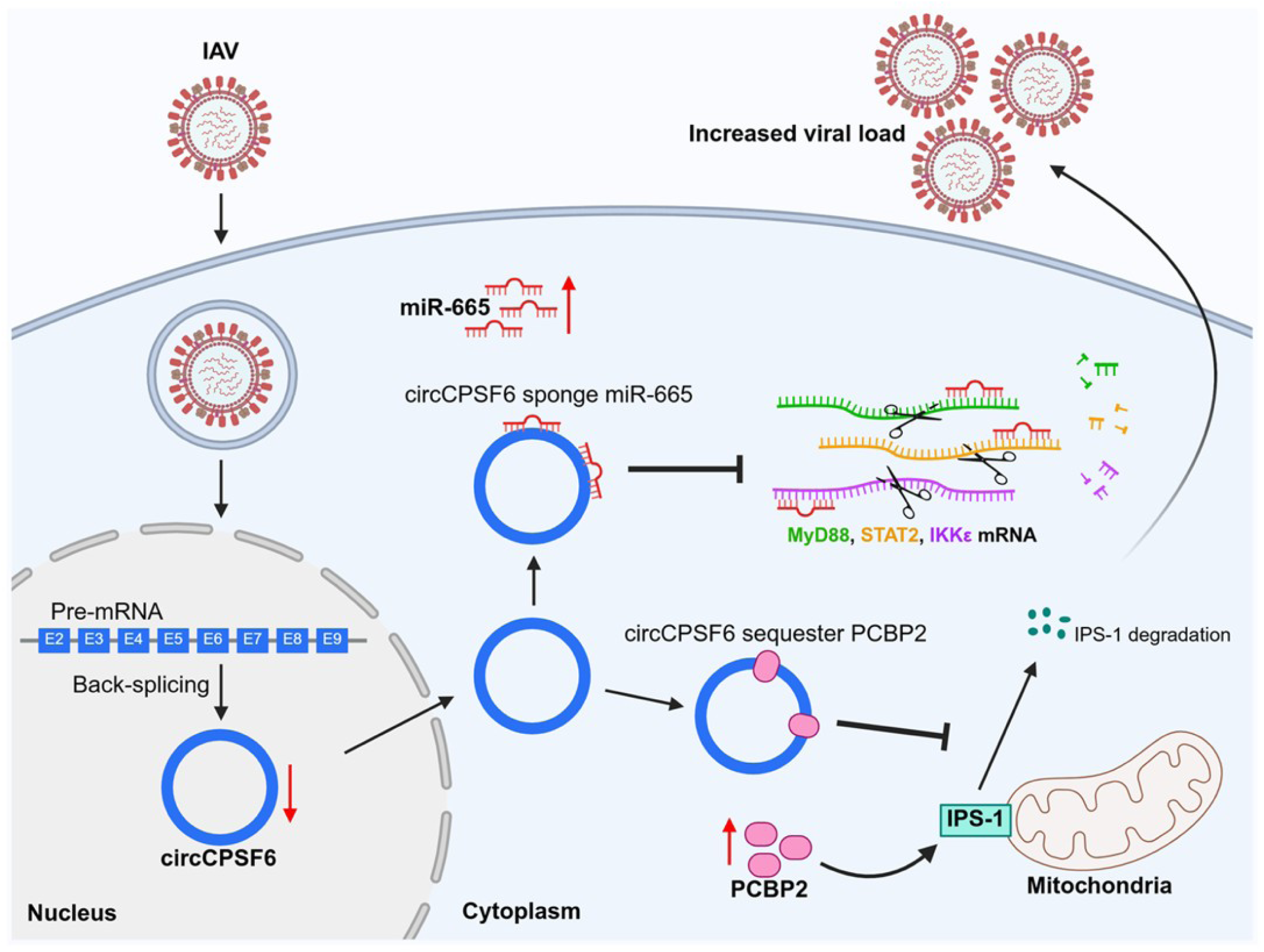

## Introduction

Influenza A virus (IAV) is a rapidly evolving respiratory pathogen, remain a recurring global health challenge due to its immune evasive capacity^1,2^. The immediate control of viral infection depends on the host innate immune system which recognize the viral genome by pattern recognition receptors (PRRs) such as cytoplasmic RIG-I like receptors (RLRs) and endosomal Toll-like receptors (TLRs), initiate the signaling cascade through specific adaptors IPS- 1/MAVS and TRIF, MYD88, respectively. These pathways converge on induction of type I and III interferons (IFN-I and -III) and proinflammatory cytokines which in turn activate the interferon-stimulated genes (ISGs), forming a critical barrier against viral replication and shaping the adaptive immunity^3,4^. A tight regulation of these pathways is critical, as insufficient response facilitate viral replication while excessive response leads to immunopathology. IAV, in turn exploit this delicate balance by subverting host regulatory network to evade the immune surveillance, highlighting the importance of comprehensive understanding of host innate antiviral mechanisms^5,6^.

Beyond the canonical protein regulators, non-coding RNAs, particularly circular RNAs (circRNAs) are increasingly appreciated as critical modulator that fine-tune antiviral signaling. CircRNAs are covalently closed, single stranded transcripts generated by back-splicing, noted for their exceptional stability and exonucleases resistance^7^. CircRNAs regulate the immune response by functioning as competing endogenous RNAs (ceRNAs) that sponges miRNAs, interact with RNA binding proteins (RBPs) or directly interact with viral proteins^8,9^. Emerging studies shows the dynamic regulation of host defense system by circRNAs during influenza infection. For example, circAIVR enhance CREBBP and IFN-β expression by sponging miR- 330-3p^10^, circMYO9A mediate SERPINE-1 driven viral restriction via sequestering miR-6059- 3p^11^, circVAMP3 decoy IAV NP and NS1 protein and restrict viral replication and pathogenesis^12^, circCBP interfere with IAV RNP assembly by interacting with NP and NS1 protein and promote stress granule dependent IFN response^13^. Despite these advances, the mechanism by which circRNAs cross-regulate both RNA-RNA and RNA-protein network to calibrate the innate immune signaling, intersection with PRR driven response remains incompletely defined.

Here, we identified circCPSF6 as a conserved circRNA which is suppressed during IAV infection, yet its presence is vital for maintaining immune homeostasis. Functional interrogation revealed circCPSF6 exert dual regulatory functions: (i) sponge the proviral miR-665, relieving repression of host antiviral genes and (ii) engage with host RBP PCBP2, antagonizing PCBP2-mediated degradation of antiviral adaptor IPS-1. Our findings reveal circCPSF6 as a central node in fine-tuning host antiviral response, broadening current understanding of non-coding RNA cross-regulation and opening avenues for antiviral therapeutics.

## Materials and Methods

### Analysis of publicly available data

For circRNA identification, high-throughput RNA-seq dataset GSE172315 (A549 cell infected with IAV H9N2 for 12h) was retrieved from NCBI-GEO repository^10^. This dataset was selected because it provides high-confidence circRNA annotation together with normalized circRNA expression value. The circRNA expression values reported as reads per millions (RPM) which is directly use for downstream analysis to visualize heatmap and bar plot using pheatmap and ggplot2 R (R 4.5.0) packages.

For miRNA identification, microarray datasets GSE103009 (A549 cell infected with IAV H1N1 for 24h)^14^ and GSE46176 (Blood sample of IAV H1N1 infected patients)^15^ were retrieved from NCBI-GEO repository and analysed in R (R 4.5.0) using limma package.

### Sequence alignment

The nucleotide sequences of human circCPSF6 and mice circCpsf6 were retrieved from CircBase database and aligned using NCBI-BLAST tool and the resulting alignment was visualized using ESPript 3.0^16^.

### Mice

This study includes CD1 (Hylasco) and C57BL/6 (NBRC, India) mice. All animal experiments were approved by the Institutional Animal Ethics Committee (IAEC) and conducted according to institutional guidelines (Approval no. 2025-IISERB-7.9-IAEC). Mice were bred and maintained under specific pathogen-free conditions at IISER Bhopal animal facility. Both male and female mice were randomly assigned to the experimental groups (n=4 per group; two males and two females).

### Cell culture

A549 human alveolar epithelial cells (ATCC CCL-185), HEK293T human embryonic kidney cells (ATCC CRL-3216), MRC5 human embryonic lung fibroblast cells (ATCC CCL-171) were maintained in Dulbecco’s Modified Eagle Medium (DMEM) supplemented with 10% Fetal Bovine Serum (FBS) and 1% penicillin-streptomycin at 37℃ in 5% CO_2_. THP1 human peripheral blood monocytes (ATCC TIB-202) were maintained in RPMI-1640 Medium supplemented with 10% Fetal Bovine Serum (FBS) and 1% penicillin-streptomycin at 37℃ in 5% CO_2_.

### PBMC and BMDC isolation

Human peripheral blood mononuclear cells (PBMCs) were isolated from healthy donors by density gradient centrifugation using Histopaque-1077 (Sigma) according to manufacturer’s instruction. Cells were maintained in RPMI-1640 Medium supplemented with 10% Fetal Bovine Serum (FBS) and 1% penicillin-streptomycin at 37℃ in 5% CO_2_.

Bone marrow–derived dendritic cells (BMDCs) were derived from wild-type (C57BL/6) mice (10-16 weeks old). Following euthanasia, hind limb was removed and bone marrow was extracted by flushing 1X PBS with 25G needle. The cells were dissociated, washed with 1X PBS at 1000g for 5 min and resuspended in DMEM supplemented with 10% Fetal Bovine Serum (FBS) and 1% penicillin-streptomycin, 25ng/mL GM-CSF. Cells were maintained at 37℃ in 5% CO_2_ and mature BMDCs were harvested after 6-7 days^17^.

### Primary lung cells isolation

Primary lung fibroblasts (mpLFs) were derived from 10-16 weeks old wild-type (C57BL/6) mice. Following euthanasia, the pulmonary circulation was perfused via right ventricle with cold 1X PBS to remove blood. The lung was harvested, finely minced and digested in cell culture media containing DNase I (0.01% w/v), collagenase and diaspase solution (TCL142, Himedia) in 1:10 ratio for 30 – 45 min at 37℃. Cell suspension were passes through 40μm strainer, washed with PBS and plated in DMEM supplemented with 10% Fetal Bovine Serum (FBS) and 1% penicillin-streptomycin at 37℃ in 5% CO_218_. The non-adherent cells were removed after 24h by PBS wash and the adherent cells were maintained in fresh cell culture medium to obtain mpLFs^19^.

### Plasmids and oligonucleotides

For overexpression of circCPSF6, the nucleotide sequence of circCPSF6 was obtained from CircBase^20^ and cloned into mc2 mNeon plasmid which was a gift from Simon Conn & Brett Stringer (Addgene Plasmid #206218). pMIR-circCPSF6 and pMIR-3’UTR plasmids were constructed by inserting full length nucleotide sequence of circCPSF6 and 3’UTRs of MyD88, STAT2, IKKε, downstream of luciferase gene in pMIR-REPORT luciferase vector, respectively. For luciferase control, pRL-TK renilla luciferase plasmid was used. pIRESneo-FLAG/HA Ago2 was a gift from Thomas Tuschl (Addgene plasmid # 10822). Full length PCBP2 was cloned into pCMV-3tag-1a vector.

CircCPSF6 BSJ specific small interfering RNA (siRNA), Cy3 labelled probe, biotin probe and PCBP2 siRNA were purchased from Eurogentec. miR-665 mimic and inhibitor were purchased from Invitrogen.

### Transfection

Plasmid DNA were transfected with jetPrime (Polyplus Transfection) reagent and siRNA, miRNA mimic/inhibitor were transfected using Lipofectamine 3000 (Invitrogen), according to manufacturer’s protocol. Human PBMCs were electroporated using the Gene Pulser Xcell electroporation system. For electroporation, 1x10^6^ PBMCs were suspended in Opti-MEM (Invitrogen) containing 50 nM siRNA/miRNA mimics, subjected to two pulses of 1000 V for 0.5 ms with a 5-second interval between pulses. Following electroporation, the cells were transferred to RPMI-1640 medium supplemented with 10% FBS.

### Virus and infection

Influenza A virus (A/PR8/H1N1) was generated as described previously^21,22^. Newcastle disease virus (NDV) and Sendai virus (SeV) were used in this study. Cells were infected with Multiplicity of Infection (MOI) 1 in serum-free DMEM for 1h, washed with 1X phosphate-buffered saline (PBS) and subsequently cultured in DMEM supplemented with 1% FBS, 1% penicillin-streptomycin and TPCK-trypsin (1μg/mL).

For *in vivo* experiment, CD1 mice (6-8 weeks old) were intranasally challenged with 100 plaque-forming units (PFU) of PR8 virus in 40μL PBS under light anaesthesia^23^. Mice were euthanised three days post infection and lungs were harvested.

### Quantitative RT-PCR

Total RNA was extracted using TRIzol reagent (Invitrogen) and cDNA was synthesized using High-Capacity cDNA Reverse Transcription Kit (Thermo Fisher Scientific) according to the manufacturer’s instructions. Gene expression was measure by quantitative real-time PCR using PowerUp SYBR Green Master Mix (Thermo Fisher Scientific) and gene specific primers **(Sup. table 3)**. For mice lung tissue samples, ∼5 mg of tissue was homogenized in a mortar and pestle and RNA was extracted using TRIzol reagent (Invitrogen). Quantification of miRNA was performed using TaqMan Fast Advanced Master Mix (Applied Biosystems) and the individual TaqMan miR-665 assay using Taqman U6 assay (Invitrogen) as a reference control.

### Digital PCR

Total RNA was extracted using TRIzol reagent (Invitrogen), cDNA was synthesized with the High-Capacity cDNA Reverse Transcription Kit (Thermo Fisher Scientific) following the manufacturer’s guidelines. Digital PCR was performed using A549 cells cDNA using gene specific primers and the QIAcuity EG PCR Kit (QIAGEN, 250111).

### RNA fluorescence *in situ* hybridization (RNA-FISH)

A549 cells were seeded on glass coverslips and infected with PR8 (MOI 1) for 24h. Cells were fixed with 4% paraformaldehyde and permeabilized with 0.1% Triton X-100. Cells were treated with prehybridization buffer (2X saline sodium citrate (SSC) buffer, 50% formamide) for 20min at room temperature. The cells were hybridized using 20ng of Cy3-labelled probe specific to back-splice junction (BSJ) of circCPSF6 in hybridization buffer (10% Dextran sulfate, 2X SSC, 50% formamide, 100 µg/mL yeast tRNA, 100 µg/mL sheared salmon sperm DNA, 1X Denhardt’s solution) at 55℃, overnight. Cells were washed with 2X SSC buffer and nuclei were stained with DAPI^24^. Images were acquired using LSM 780 confocal laser microscope.

### Immunoblotting analysis

Cells were harvested 24h post infection, washed with PBS and lysed in ice-cold lysis buffer containing 1X protease inhibitor cocktail (11836145001, Roche). Cell lysates were centrifuged at 15000g for 15min at 4℃ and supernatants were collected. Protein was quantified by Bradford assay and ∼20μg of protein per sample was resolved by SDS–PAGE. Immunoblotting was performed using antibodies against IAV-NP (PA5-32242, Invitrogen), MYD88 (sc-74532, Santa Cruz), STAT2 (sc-514193, Santa Cruz), IKBKE (PA5-87387, Invitrogen), FLAG (F1804, Sigma-Aldrich), PCBP2 (PA5-30116, Invitrogen), IPS-1 (sc-166583, Santa Cruz) and β-actin (A1978, Sigma-Aldrich). Secondary anti-mouse and anti-rabbit IgG antibodies were purchased from Invitrogen, and blots were visualized using the LI-COR imaging system.

### Immunofluorescence assay

A549 cells were seeded on glass coverslips, transfected with siRNA or mimic for 24h and infected with PR8 (MOI 1) for 24h. Cells were fixed with 4% paraformaldehyde, permeabilized with 0.1% Triton X-100 and incubated with 5% FBS containing blocking solution for 1h at room temperature. Cells were incubated with primary antibodies and appropriate fluorophore-conjugated secondary antibodies. Cell nuclei were counterstained with DAPI and images were acquired with LSM 780 confocal laser microscope. For colocalization analysis of circCPSF6 and PCBP2, A549 cells were subjected to immunostaining of PCBP2. Briefly, cells were washed, fixed with fresh 4% paraformaldehyde for 5min at room temperature and subsequently RNA-FISH was performed using Cy3 labelled circCPSF6 probe^24^. Colocalization signal was quantified by line-scan intensity profiling (ImageJ/Fiji).

### Flow cytometry

A549 cells were harvested 24h post infection, trypsinized, fixed with 4% paraformaldehyde and permeabilised with 0.1% Triton X-100 for 10min. Cells were incubated with 5% FBS containing blocking solution for 45min and incubated with primary antibody for 1h followed by secondary antibody for 30min. Cells were analysed on FACSAria III flow cytometer (Becton Dickinson) and data were analysed using FlowJo software.

### RNA immunoprecipitation (RIP) assay

The Ago2-RIP assay was performed in HEK293T cells after co-transfection of Flag-Ago2 and miR-665 mimic, followed by PR8 infection. After 24h infection, cells were lysed in ice-cold lysis buffer supplemented with 1X protease inhibitor cocktail (11836145001, Roche) and lysate supernatant was collected. The clarified lysate was incubated with FLAG M2 affinity beads (Sigma) at 4℃ overnight. Beads were washed thoroughly; bound RNA was eluted using TRIzol reagent and subjected to RT-PCR and normalized with 10% input of indicated RNA.

### Luciferase reporter assay

HEK293T cells were seeded in 24 well plate and co-transfected with 100ng of luciferase reporter plasmid, 20ng of *Renilla* luciferase plasmid, 25nM of control/miR-665 mimic or 50nM of inhibitor control/miR-665 inhibitor. Cells were collected and lysed after 36h post transfection and luciferase activity was quantified using Dual-Luciferase Reporter Assay System (E1960, Promega) according to the manufacturer’s instructions and the readings were recorded in the GloMax Multi+ system (Promega).

### RNA pulldown assay

Uninfected or PR8 infected A549 or HEK293T cells were harvested and lysed in ice-cold lysis buffer supplemented with 1X protease inhibitor cocktail (11836145001, Roche). Lysate were clarified by centrifugation and 10% of lysate was aliquoted for input. Pierce Streptavidin Magnetic Beads (88817, Thermo Fisher Scientific) were pre-incubated with circCPSF6 BSJ specific biotin-labelled probe or control probe for 2h at room temperature followed by overnight incubation with remaining clarified lysate at 4℃. The beads were washed with ice-cold lysis buffer and bound RNA was eluted using TRIzol reagent. For protein pulldown, the beads were washed with ice-cold lysis buffer and bound proteins were eluted in 50μL elution buffer by heating at 100℃ for 5min^25^.

### Mass spectrometry

Protein eluted by RNA pulldown assay were resolved by 12% SDS-PAGE gel. When protein bands entered separating gel, electrophoresis was stopped to stack the protein. The protein bands were excised, destained and subjected to in-gel digestion, including reduction (10mM DTT + 100mM Ammonium bicarbonate (ABC)) and alkylation (50mM iodoacetamide (IAA) + 100mM ABC), followed by overnight trypsin (1ng/mL; Promega) digestion at 37℃. Tryptic peptides were extracted using extraction solution (0.1% Trifluoroaceticacid (TFA) + 50% Acetonitrile), sonicated for 10min in water bath, pooled and dried in SpeedVac concentrator^26^. Dried peptide samples were subjected to the Mass Spectrometry. Raw spectral data were processed using MaxQuant (v2.7.3.0)^27^ with default parameters and Human UniProt FASTA database (May 2024) for protein identification and quantification. The output data of MaxQuant was analysed in Perseus (v2.1.6.0), where contaminants, reverse hits, and proteins identified only by site were excluded. Protein intensity was log2 transformed, rows with at least 70% valid values across replicates were retained, and missing values were imputed from a normal distribution using default Perseus settings. Annotation and matrix normalization (z-score) were performed for comparative visualization as replicate and group mean heatmaps^28^.

### Enzyme-linked immunosorbent assay (ELISA)

Culture supernatants from A549 cells and PBMCs were collected and analysed by ELISA following the manufacturer’s protocols to determine the levels of IP-10 (550926, BD OptEIA Human IP-10 ELISA Set) and TNF-α (555212, BD OptEIA Human TNF-α ELISA Set).

### Statistical analysis

All experiments included appropriate control or mock-transfected samples and were independently repeated two to three times. Data were shown as mean ± SEM. Data were analysed using GraphPad Prism (v8.0.2). Comparison between two groups were performed with unpaired, two-tailed Student’s *t*-test whereas multiple group comparisons were evaluated with two-way ANOVA. *P* value < 0.05 were considered statistically significant. In figures, statistical significance is denoted as follows: ****, *p* < 0.0001; ***, *p* < 0.001; **, *p* < 0.01; *, *p* < 0.05; ns, not significant.

## Results

### Identification and characterization of circCPSF6

To investigate the circRNA dynamics during influenza virus infection, we analysed publicly available total RNA-sequencing data from GEO database (GSE172315) which profiles IAV H9N2 infected human lung epithelial cell, A549 at 12h post infection. The dataset provides normalized circRNA expression value in reads per million (RPM) focusing on reads spanning unique back-spliced junctions (BSJs) revealing numerous dysregulated circRNAs **(Fig. 1A)**. Among these, the expression level of circCPSF6 (hsa_circ_0000417) was consistently reduced in infected sample compare to control **(Fig.1B)**. Sequence conservation analysis further determined strong evolutionary conservation (92%) of circCPSF6 in mice **(Fig. S1A)**.

**Fig 1.**
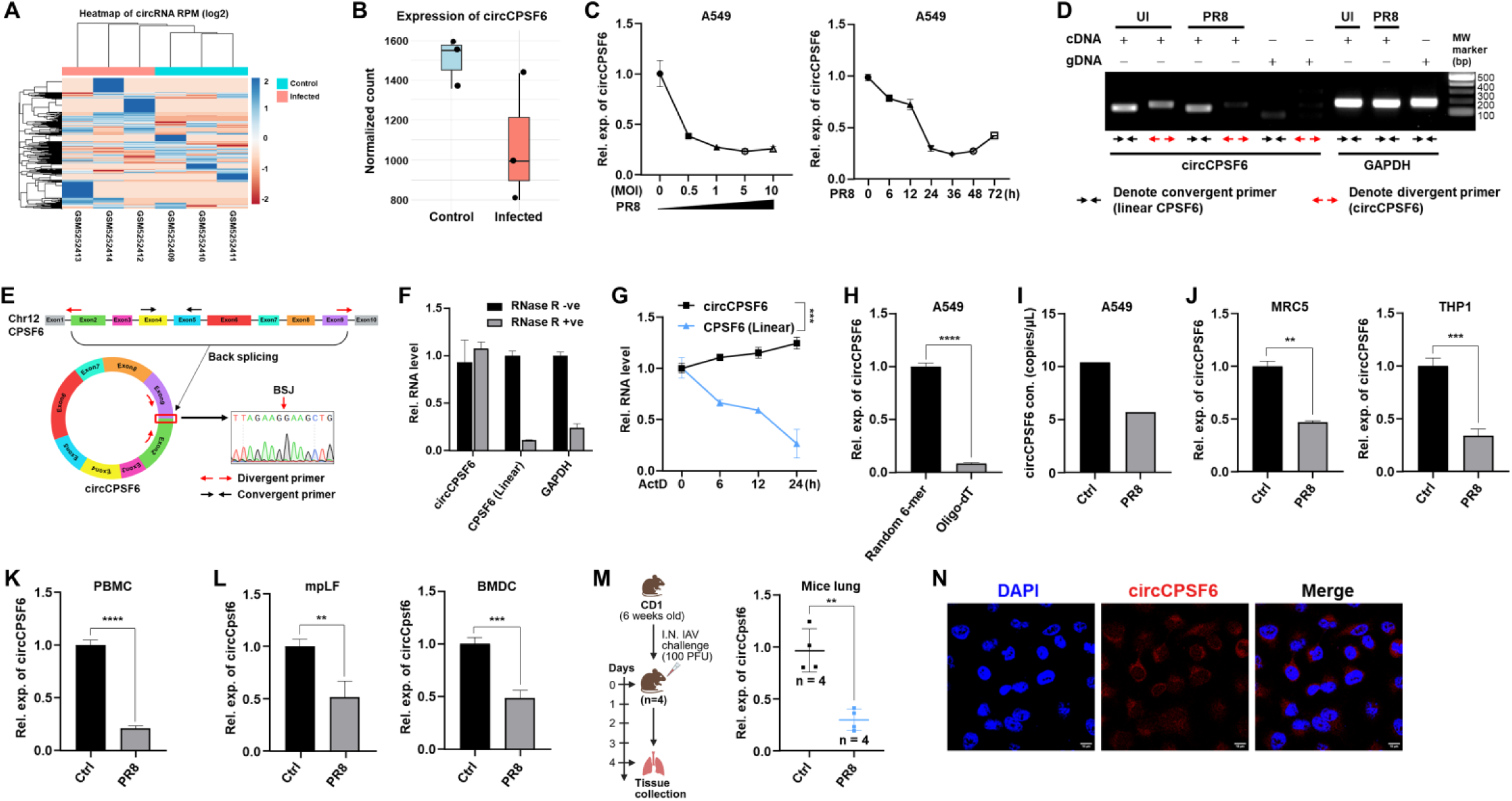
Identification and characterization of circCPSF6 during IAV infection. **(A)** Heatmap showing clustering of log2 transformed reads per million (RPM) value of circRNAs from RNA seq data (GSE172315) in IAV H9N2 infected A549 cells with relative expression values indicated by blue to red scale (blue, high; red, low). **(B)** CircCPSF6 expression in control and infected sample from same data. **(C)** RT-PCR validation of circCPSF6 expression level in A549 cells, infected with PR8 at indicated multiplicity of infection (MOI) and post infection time points. **(D)** PCR amplification of circCPSF6 and linear CPSF6 using divergent (circRNA specific) and convergent (linear RNA) primers in cDNA or genomic DNA (gDNA) from A549 cells; GAPDH used as control. **(E)** Schematic representation of circCPSF6 genomic location generated by the back splicing of CPSF6 exons and Sanger sequencing of divergent primer amplicon showing back splice junction (BSJ). **(F)** Expression of circCPSF6, linear CPSF6 and GAPDH measured with or without RNase R (2U/μg RNA) treatment at 37℃ for 20 mins in A549 cells. **(G)** Expression of circCPSF6 and linear CPSF6 measured in actinomycin D (2μg/mL) treated A549 cells for indicated time. **(H)** CircCPSF6 amplification from cDNA prepared using random hexamer and oligo-dT primer. **(I)** CircCPSF6 copy number quantified by digital PCR (dPCR) in PR8 infected (MOI 1; 24h) A549 cells. **(J – L)** CircCPSF6 expression in human cell lines **(J)** MRC5 and THP1, **(K)** human primary PBMCs, **(L)** murine primary lung fibroblasts (mpLF) and BMDCs, post PR8 infection (MOI 1; 24h). **(M)** Schematic representation of *in vivo* virus challenge and expression of circCpsf6 in mice lung tissue after PR8 infection (PFU 100; 3 days). **(N)** RNA-FISH using Cy3 labelled antisense-probe designed specific to circCPSF6 BSJ (red) for the cellular localization. Nuclei were stained with DAPI (blue). Scale bar, 10μm. Data are presented as the mean ± SEM from triplicate samples of a single experiment and representative of three independent experiments. **** *p* < 0.0001, *** *p* < 0.001, ** *p* < 0.01 by two-tailed unpaired Student’s *t*-test or two-way ANOVA test. (Schematics are created with BioRender.com)

Experimental validation in A549 human lung epithelial cells using cDNA prepared by random hexamer primer confirmed significant downregulation of circCPSF6 upon IAV PR8 infection in dose and time dependent manner **(Fig. 1C)**. To verify the circular nature of circCPSF6, divergent and convergent primers were designed for circCPSF6 and linear CPSF6, yielding 144bp and 120bp amplicon, respectively. Both amplicons were detected from cDNA, whereas the divergent primer failed to amplify from genomic DNA **(Fig. 1D)**, confirming specific back-splicing. Sanger sequencing of divergent primer amplicon verified the back-splice junction (BSJ) connecting exon 2 and 9 of CPSF6 on chromosome 12 **(Fig. 1E)**, validating the circular structure of circCPSF6. RNase R digestion and actinomycin D transcriptional blockade assays further validated the hallmark stability of circCPSF6 compared to its linear counterpart **(Fig. 1F, G)**, while selective amplification from cDNA prepared using random hexamers but not oligo-dT primers, confirmed the non-polyadenylated circular structure **(Fig. 1H)**.

Further, absolute quantification by digital PCR (dPCR) affirmed reduced copy number of circCPSF6 in PR8 infected A549 cells **(Fig. 1I)**. Similar downregulation was observed in other human cell types, including MRC5, THP1 and human primary PBMCs, infected with PR8 **(Fig. 1J, K)**. Consistent with its conservation, circCPSF6 expression was also downregulated in PR8 infected mice primary lung fibroblasts (mpLFs) and mice bone marrow-derived dendritic cells (BMDCs) **(Fig. 1L)**. *In vivo* PR8 infection in wild type CD1 mice (as shown in schematic) corroborate significant decrease in circCPSF6 expression in lung tissue **(Fig. 1M)**. Importantly, suppression of circCPSF6 was not limited to IAV, as infection with Newcastle disease virus (NDV) also led to reduced expression in A549 and MRC5 cells **(Fig. S1B, S1C)**. Additionally, RNA-FISH using Cy3-labelled probe targeting the BSJ revealed predominant cytoplasmic localization of circCPSF6 in A549 cells **(Fig. 1N)**.

Collectively, all these results indicate that circCPSF6 is a conserved circRNA, broadly suppressed *in vitro* and *in vivo* model during RNA virus infections and predominantly present in cell cytoplasm.

### CircCPSF6 restricts IAV replication through inflammatory signaling

To investigate the biological function of circCPSF6 during IAV infection, siRNA was designed specifically targeting the BSJ of circCPSF6 without affecting the linear CPSF6 transcript **(Fig. 2A)**. Knockdown of circCPSF6 significantly enhanced PR8 burden in A549 cells, as well as in MRC5 cells, as measured by RT-PCR **(Fig. 2B, S2A)**. Consistent results were obtained in primary cells including human PBMCs and murine mpLFs and BMDCs upon circCPSF6 knockdown **(Fig. 2C, D, S2B)**. Elevated viral load was also detected in other RNA virus infections such as NDV and Sendai virus (SeV) following circCPSF6 knockdown in A549 cells **(Fig. S2C, S2D)**. Further, we confirmed increased viral replication at protein level by immunoblotting in A549 and mpLF cells **(Fig. 2E)**. Consistently, flow cytometry and immunofluorescence also demonstrated enhanced viral protein in A549 cells **(Fig. 2F, G)**.

**Fig 2.**
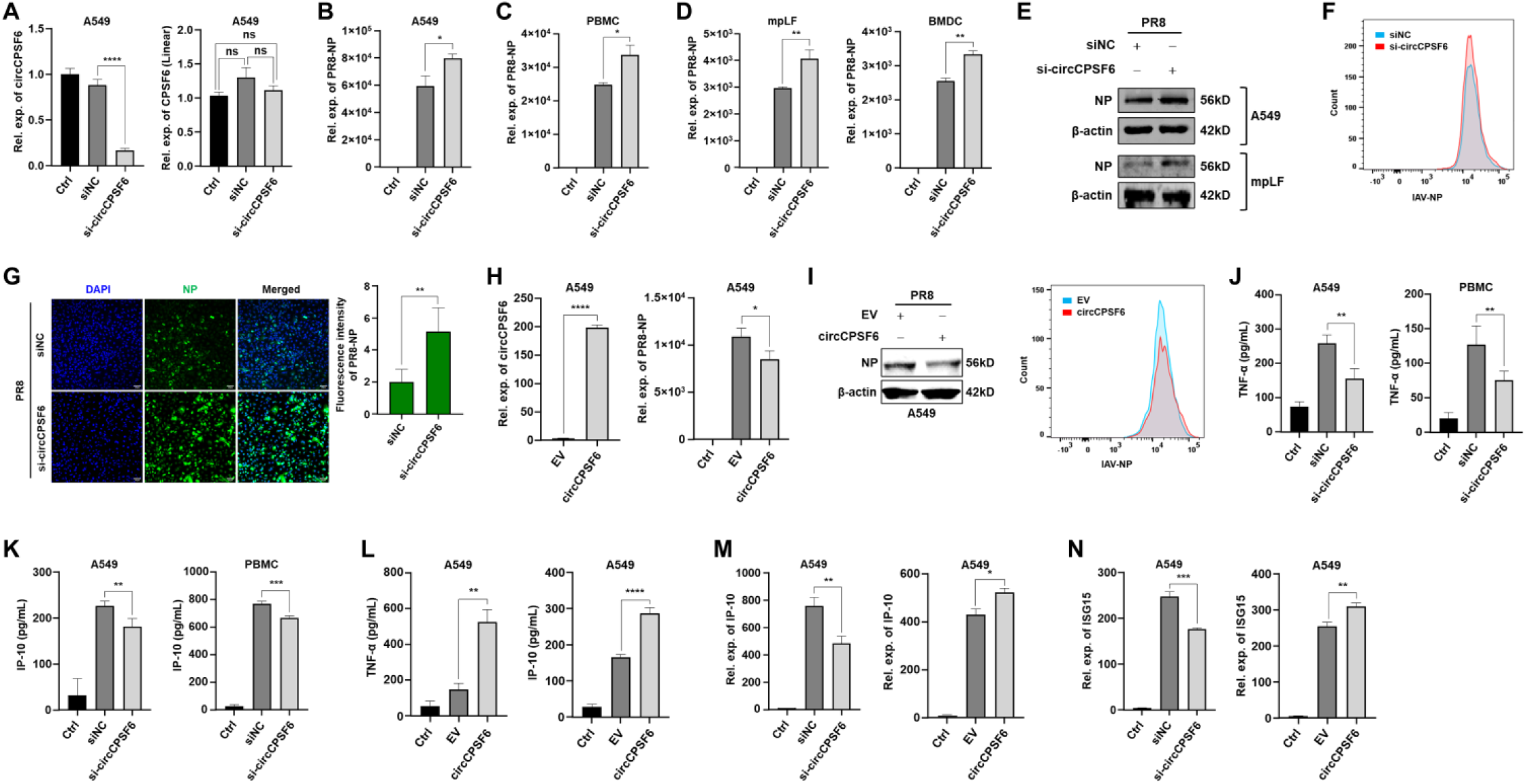
circCPSF6 inhibit IAV replication and induce inflammatory response. **(A)** BSJ specific siRNA mediated knockdown of circCPSF6 and expression level of linear CPSF6 was determined by RT-PCR. **(B)** Viral load determined in circCPSF6 knockdown, PR8 infected (MOI 1; 24h) A549 cells by RT-PCR. **(C – D)** Viral load determined by RT-PCR in circCPSF6 knockdown, **(C)** human PBMCs, **(D)** mice mpLFs and BMDCs after PR8 infection (MOI 1; 24h). **(E)** Viral protein expression determined by immunoblotting in circCPSF6 knockdown, PR8 infected (MOI 1; 24h) A549 cells and mpLFs. **(F)** Flow cytometry analysis with IAV NP antibody in circCPSF6 knockdown, PR8 infected (MOI 1; 24h) A549 cells. **(G)** Immunofluorescence with IAV NP antibody (green) in circCPSF6 knockdown, PR8 infected (MOI 1; 24h) A549 cells. Nuclei stained with DAPI. Scale bar, 50μm. The graph showing the fluorescence intensity of IAV NP protein. **(H)** Overexpression of circCPSF6 and viral load measured by RT-PCR in PR8 infected (MOI 1; 24h) A549 cells. **(I)** Immunoblot and flow cytometry analysis showing IAV NP protein level in circCPSF6 overexpressing A549 cells following PR8 infection (MOI 1; 24h). **(J – K)** Quantification of cytokines **(J)** TNF-α and **(K)** IP-10 by ELISA in culture supernatant of circCPSF6 knockdown A549 cells and human PBMCs, infected with PR8 (MOI 1; 24h). **(L)** Quantification of cytokines TNF-α and IP-10 by ELISA in culture supernatant of circCPSF6 overexpressing A549 cells, followed by PR8 infection (MOI 1; 24h). **(M)** IP-10 transcript expression measured by RT-PCR in circCPSF6 knockdown or overexpressing A549 cells following PR8 infection (MOI 1; 24h). **(N)** Transcript expression of ISG15 in circCPSF6 knockdown or overexpressing A549 cells followed by PR8 infection (MOI 1; 24h). Data are presented as the mean ± SEM from triplicate samples of a single experiment and representative of three independent experiments. **** *p* < 0.0001, *** *p* < 0.001, ** *p* < 0.01, * *p* < 0.05, ns is non-significant by two-tailed unpaired Student’s *t*-test.

Conversely, ectopic expression of circCPSF6 significantly restrained PR8 replication in both A549 and MRC5 cells **(Fig. 2H, S2E)**. Additionally, immunoblotting and flow cytometry **(Fig. 2I)** results confirmed reduced viral protein expression upon circCPSF6 overexpression in A549 cells. Importantly, circCPSF6 overexpression also curtailed NDV infection in A549 cells **(Fig. S2F)**, implicating a broad antiviral response.

Next, we examined cytokine response in culture supernatants by ELISA, where circCPSF6 knockdown followed by PR8 infection in A549 cells and human PBMCs resulted suppressed TNF-α and IP-10 secretion **(Fig. 2J, K)**. Conversely, ectopic expression of circCPSF6 in A549 cells followed by PR8 infection, induced TNF-α and IP-10 secretion **(Fig. 2L)**. Consistently, transcript level expression of IP-10 showed similar result with ELISA in A549 cells upon knockdown and overexpression of circCPSF6 **(Fig. 2M)**. Notably, IL-6 transcript was also decreased in circCPSF6 knockdown, PR8 infected A549 cells **(Fig. S2G)**. Moreover, the antiviral effector, ISG15 was significantly suppressed following circCPSF6 knockdown whereas upregulated with circCPSF6 overexpression, as determined by RT-PCR **(Fig. 2N)**.

Altogether, these results suggest circCPSF6 expression linked with viral replication and induction of antiviral innate cytokines, hence, contributing to regulation of host antiviral defence during viral infection.

### CircCPSF6 sponges miR-665, a virus induced miRNA

By validating circCPSF6 is downregulated and act as an antiviral regulator during IAV infection, we further explored the underlying molecular mechanism. One of the key functions of circRNA in physiological state is to function as molecular sponge for miRNA thereby regulate gene expression post transcriptionally, hence, we sought to identify miRNAs interacting with circCPSF6. Circinteractome^29^ search revealed several miRNAs with putative binding sites on circCPSF6 **(Fig. 3A)**. To prioritize miRNAs functionally relevant to IAV infection, we examined two independent datasets (GSE103009 and GSE46176) from GEO database for significantly dysregulated miRNAs (adj. *P* value < 0.05) and intersection of these miRNAs with Circinteractome candidates highlighted a single shared miRNA, hsa-miR-665 (miR-665) which function is not explored in context of viral infection **(Fig. 3B, S3A)**. Moreover, the binding affinity prediction using miRanda algorithm revealed multiple high affinity binding sites for miR-665 that are conserved between human and mice circCPSF6 **(Sup. table 1, 2)**, illustrated schematically in **Fig. 3C**.

**Fig 3.**
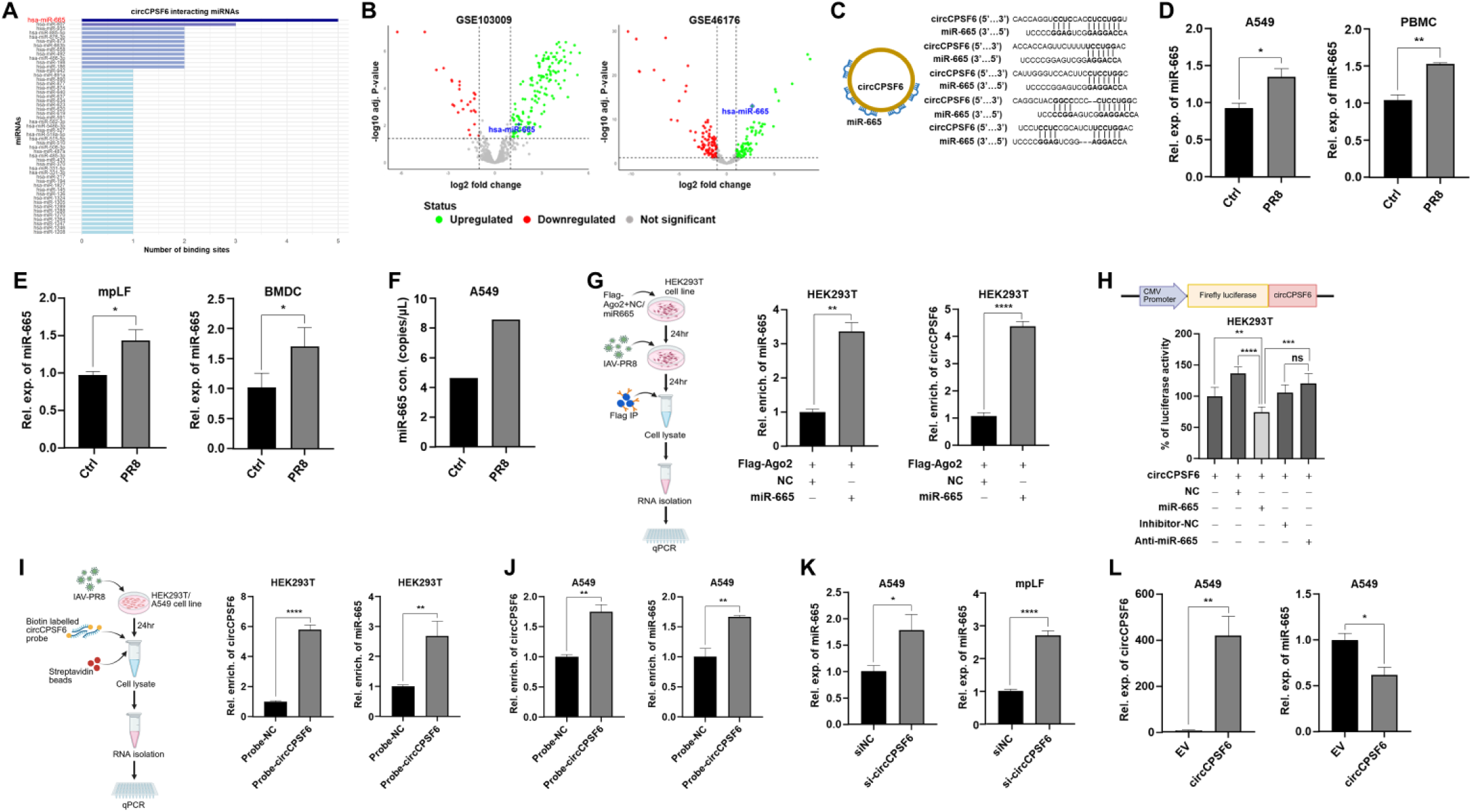
miR-665 sponged by circCPSF6. **(A)** CircCPSF6 interacting miRNAs identified from Circinteractome database. **(B)** Volcano plot of differentially expressed miRNAs from two independent GEO datasets, GSE103009 and GSE46176 comparing influenza virus infected and control samples. Each point represents individual miRNA (green, upregulated; red, downregulated; log_2_ fold change > 1; adjusted *P* value > 0.05). In both datasets, miR-665 (blue diamond) demonstrating induction during infection. **(C)** Schematic representation of miR-665 binding sites on circCPSF6. **(D)** Expression level of miR-665 measured by RT-PCR in A549 cells and human PBMCs following PR8 infection (MOI 1; 24h). **(E)** miR-665 expression in PR8 infected (MOI 1; 24h) mice mpLFs and BMDCs measured by RT-PCR. **(F)** Copy number of miR-665 determined by dPCR in PR8 infected (MOI 1; 24h) A549 cells. **(G)** Schematic representing workflow for RIP assay. Enrichment of miR-665 and circCPSF6 measured by RT- PCR, relative to respective input samples. **(H)** Schematic illustrating luciferase reporter construct containing circCPSF6 sequence. Luciferase assay demonstrating percentage of luciferase activity in HEK293T cells co-transfected with luciferase reporter and mimic NC/mimic miR-665, miRNA inhibitor NC/miR-665 inhibitor (Anti-miR-665). **(I - J)** Schematic illustrating workflow for RNA pulldown assay with circCPSF6 BSJ specific biotin labelled probe. Enrichment of circCPSF6 and miR-665 measured in **(I)** HEK293T and **(J)** A549 cells by RT-PCR, relative to respective input samples. **(K)** Expression of miR-665 measured by RT-PCR in circCPSF6 knockdown A549 cells and mpLFs following PR8 infection (MOI 1; 24h). **(L)** Expression level of circCPSF6 and miR-665 measured in circCPSF6 overexpressing A549 cells following PR8 infection (MOI 1; 24h) by RT-PCR. Data are presented as the mean ± SEM from triplicate samples of a single experiment and representative of three independent experiments. **** *p* < 0.0001, *** *p* < 0.001, ** *p* < 0.01, * *p* < 0.05, ns is non-significant by two-tailed unpaired Student’s *t*-test. (Schematics are created with BioRender.com)

Corroborating with the bioinformatics prediction, miR-665 expression was significantly induced in PR8 infected A549 cells and human PBMCs as well as in murine mpLFs and BMDCs **(Fig. 3D, E)**. Similar induction of miR-665 was observed in NDV infected A549 cells **(Fig. S3B)**. Absolute quantification by dPCR determined increased copy number of miR-665 in PR8 infected A549 cells **(Fig. 3F)**.

We next validated the direct interaction between circCPSF6 and miR-665 by RNA immunoprecipitation (RIP) assay with Flag-tagged Ago2 protein (as shown in schematic), a core component of RNA induced silencing complex (RISC). This resulted robust enrichment of miR-665 and circCPSF6 in Ago2 immunoprecipitate, especially with miR-665 overexpression fraction, demonstrating their association with RISC **(Fig. 3G)**. This interaction was functionally validated by luciferase reporter assay demonstrating co-expression of miR- 665 with circCPSF6 containing reporter, significantly suppress luciferase activity, whereas co-expression with miR-665 specific antisense inhibitor reversed this effect **(Fig. 3H)**. Consistently, RNA pulldown assay using biotinylated probe specific to circCPSF6 BSJ (as shown in schematic), confirmed robust enrichment of endogenous circCPSF6 and miR-665, compare to negative control-probe (probe-NC), in PR8 infected HEK293T and A549 cells **(Fig. 3I, J)**.

Additionally, functional interplay between circCPSF6 and miR-665 was analysed where circCPSF6 knockdown increased miR-665 expression in PR8 infected A549 and mpLF cells **(Fig. 3K, S3C)**, in contrast, overexpression of circCPSF6 led to decreased miR-665 expression in PR8 infected A549 cells **(Fig. 3L)**.

Together all these results provide comprehensive evidence that circCPSF6 interact and sponge miR-665, a virus-induced novel host miRNA, and modulate its expression, suggesting a regulatory mechanism that may shape the host innate antiviral response.

### miR-665 enhance influenza virus replication

Based on our preceding results demonstrating circCPSF6 act as a molecular sponge for miR- 665 and it is upregulated during virus infection, we sought to delineate the functional role of miR-665 during viral infection. Overexpression of miR-665 in A549 cells and human PBMCs followed by PR8 infection resulted significantly increased viral load **(Fig. 4A, B)**, a result recapitulated in murine mpLFs and BMDCs infected with PR8 **(Fig. 4C, D)**, underscoring its conserved role across species. To determine whether the proviral activity of miR-665 extend to other RNA virus, we overexpressed miR-665 in A549 cells and infected with NDV and SeV. In both cases, viral load was significantly enhanced upon miR-665 overexpression **(Fig. 4SA, 4SB)**.

**Fig 4.**
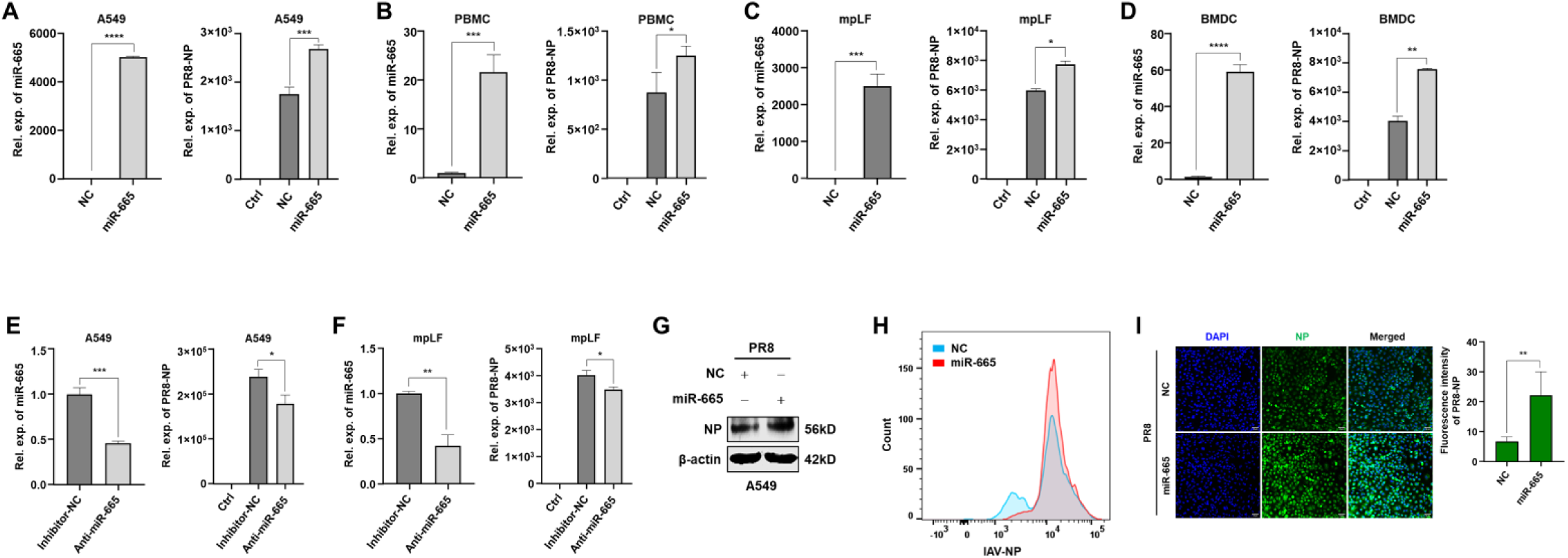
miR-665 enhance influenza virus replication. (A -. **B)** Overexpression level of miR- 665 and viral load measured by RT-PCR in PR8 infected (MOI 1; 24h) **(A)** A549 cells and **(B)** human PBMCs. **(C - D)** Overexpression level of miR-665 and viral load measured by RT-PCR in PR8 infected (MOI 1; 24h) mice **(C)** mpLFs and **(D)** BMDCs. **(E - F)** Expression level of miR-665 and viral load measured by RT-PCR after silencing of miR-665 by antisense inhibitor, followed by PR8 infection (MOI 1; 24h) in **(E)** A549 cells and **(F)** mpLFs. **(G)** Viral protein determined by immunoblotting in miR-665 overexpressing A549 cells followed by PR8 infection (MOI 1; 24h). **(H)** Flow cytometry analysis with IAV NP antibody in miR-665 overexpressing A549 cells followed by PR8 infection (MOI 1; 24h). **(I)** Immunofluorescence with IAV NP antibody (green) in miR-665 overexpressing, PR8 infected (MOI 1; 24h) A549 cells. Nuclei stained with DAPI. Scale bar, 50μm. The graph showing the fluorescence intensity of IAV NP protein. Data are presented as the mean ± SEM from triplicate samples of a single experiment and representative of three independent experiments. **** *p* < 0.0001, *** *p* < 0.001, ** *p* < 0.01, * *p* < 0.05 by two-tailed unpaired Student’s *t*-test.

Conversely, targeted sequestering of endogenous miR-665 with specific antisense inhibitor significantly reduced the viral burden in PR8 infected A549 cells and mpLFs **(Fig. 4E, F)**. Further, the analysis of viral protein expression by immunoblotting revealed increased abundance upon miR-665 overexpression in PR8 infected A549 cells **(Fig. 4G)**. Flow cytometry and immunofluorescence analysis corroborated these findings with a higher proportion of viral antigen-positive cells in miR-665 overexpressing PR8 infected A549 cells **(Fig. 4H, I)**.

Collectively, all these results indicate miR-665 as a proviral factor during influenza virus as well as other RNA virus infection, suggesting that circCPSF6 mediated sequestration of miR- 665 exert the antiviral function.

### miR-665 targets host genes pivotal for the antiviral signalling pathways

Given that miRNAs regulate post-transcriptional gene expression by binding to target sites within the 3’UTR of mRNAs, we investigated potential genes targeted by miR-665. Integrative analysis of miRDB and ENCORI predictions yielded 335 overlapping genes, and the pathway enrichment analysis revealed enrichment of influenza A virus infection pathway along with other viral and immune related pathways, emphasizing the potential involvement of miR-665 target genes in antiviral response **(Fig. 5A)**. Among these, MyD88, STAT2 and IKKε were selected for further study due to their central role in antiviral signaling cascades. To validate the interaction of these candidate targets with miR-665, luciferase reporter assay was performed which demonstrated co-transfection of miR-665 with reporter containing 3’UTRs of MyD88, STAT2 or IKKε led to reduction in luciferase activity significantly, whereas co-transfection with miR-665 inhibitor reversed this effect **(Fig. 5B)**. Consistently, RIP assay using Flag-Ago2 resulted marked enrichment of MyD88, STAT2 and IKKε transcripts in miR-665 co-transfected fraction, further supporting their recruitment to miR-665-associated RISC and bona fide interactions (**Fig. 5C)**. To assess regulation of these antiviral genes at protein level, we performed immunoblotting in A549 and HEK293T cells overexpressing either miR-665 or circCPSF6 followed by PR8 infection. Overexpression of miR-665 resulted in decreased protein levels of MyD88, STAT2 and IKKε, whereas circCPSF6 overexpression restored their expression **(Fig. 5D, E)**.

**Fig 5.**
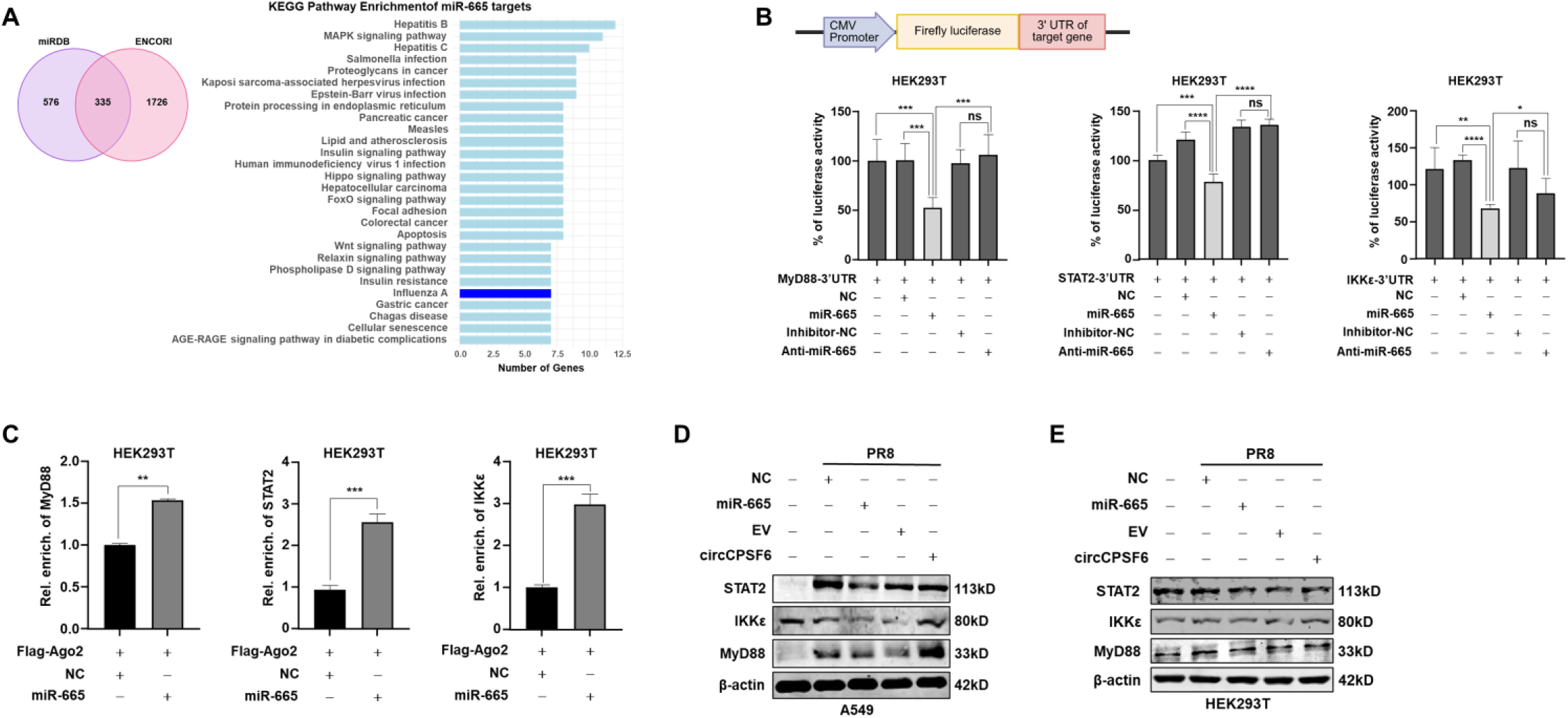
miR-665 targets host anti-viral genes. **(A)** Venn diagram showing overlap of miR- 665 host target genes among miRDB and ENCORI databases. Bar graph indicating enriched KEGG pathways among the common genes, analysed by pathway analysis. Blue bar showing number of miR-665 host target genes involved in influenza A infection pathway. **(B)** Schematic illustrating luciferase reporter construct containing 3’UTRs of miR-665 targets, MyD88, STAT2 and IKKε. Luciferase assay demonstrating luciferase activity in HEK293T cells co-transfected with respective luciferase reporter and mimic NC/mimic miR-665, miRNA inhibitor NC/miR-665 inhibitor (Anti-miR-665). **(C)** RIP assay with Flag-Ago2 along with NC/miR-665, following PR8 infection (MOI 1; 24h) in HEK293T cells, showing enrichment of MyD88, STAT2 and IKKε, relative to respective input samples, measured by RT-PCR. **(D – E)** Immunoblotting of MyD88, STAT2 and IKKε in **(D)** A549 and **(E)** HEK293T cells after overexpressing miR-665 or circCPSF6 followed by PR8 infection (MOI 1; 24h). Data are presented as the mean ± SEM from triplicate samples of a single experiment and representative of three independent experiments. **** *p* < 0.0001, *** *p* < 0.001, ** *p* < 0.01, * *p* < 0.05, ns is non-significant by two-tailed unpaired Student’s *t*-test. (Schematic is created with BioRender.com)

These findings indicate that miR-665 interact and suppress pivotal antiviral mediators, while circCPSF6 can antagonize this suppression by sequestering miR-665, suggesting a dynamic immune regulation. Thus, highlighting the important role of circCPSF6 in maintaining antiviral immune response.

### Identification of circCPSF6 interacting proteins during IAV infection

Although circRNAs are established as miRNA sponges, the contribution of circRNA–RNA binding protein interactions to gene regulation during viral infection remains insufficiently characterized. Therefore, to further elucidate the function of circCPSF6, we performed RNA pull down using biotinylated probe specific to BSJ of circCPSF6 in PR8 infected A549 cells followed by mass spectrometry as shown in the schematic **(Fig. 6A)**. The mass spectrometry analysis identified more than 100 proteins enriched with circCPSF6 **(Fig. S5A)** with PCA plot demonstrating clear segregation of sample groups **(Fig. S5B)**. To establish high confidence interactor of circCPSF6, statistical filtering and enrichment analysis using Perseus tool^30^ reported 49 proteins **(Fig. 6B, S5C)**. Among these, PCBP2 protein was prioritise as it is consistently enriched and has established role in viral infection and PRR-mediated innate immune response. To confirm the interaction between circCPSF6 and PCBP2, we performed RNA pulldown assay using circCPSF6 BSJ specific biotinylated probe, which revealed enrichment of PCBP2 by immunoblotting in both uninfected and modestly enhanced in PR8- infected A549 cells **(Fig. 6C, S5D)**. Complementary RIP assay with Flag-tagged PCBP2 resulted highly enriched circCPSF6 in contrast to the empty vector fraction in PR8 infected HEK293T cells **(Fig. 6D)**. Additionally, combined RNA-FISH of circCPSF6 and immunostaining of PCBP2 revealed predominant cytoplasmic co-localization of circCPSF6 and PCBP2 in A549 cells, corroborated by overlapping fluorescence intensity profiles across measured distance **(Fig. 6E)**. All these results indicate that circCPSF6 interact with PCBP2 in the cell cytoplasm, suggesting a potential regulatory axis that may influence host antiviral response during influenza virus infection.

**Fig 6.**
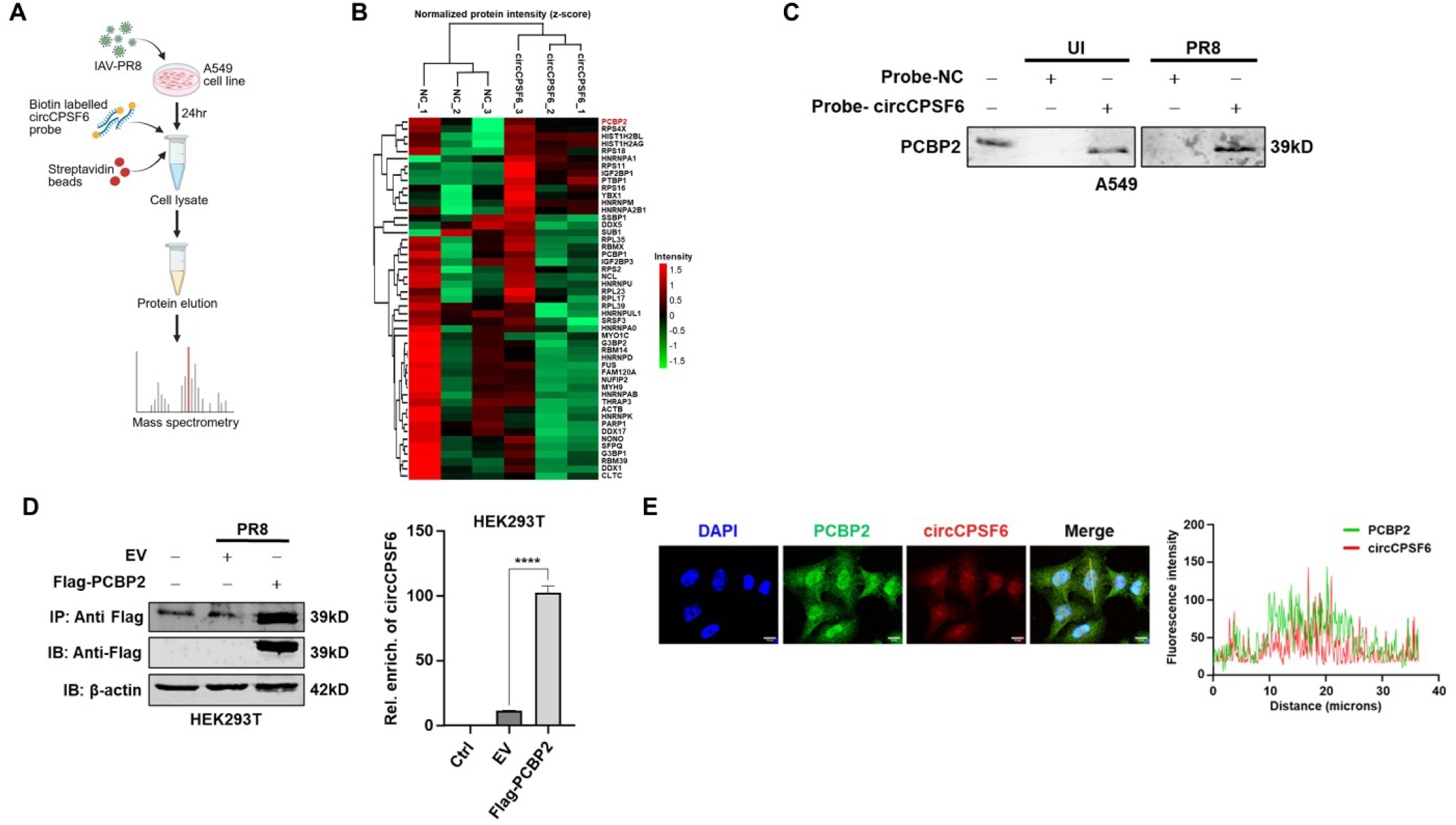
Identification of circCPSF6 interacting proteins. **(A)** Schematic representing workflow of RNA pulldown assay with circCPSF6 BSJ specific biotin labelled probe following mass spectrometry to identify circCPSF6 interacting proteins. **(B)** Heatmap showing clustering of normalized protein intensity (Z-score) from mass spectrometry data with relative enrichment showing in red and green scale (red, high; green, low). **(C)** RNA pulldown assay with biotin labelled probe specific to circCPSF6 BSJ in A549 cells with or without PR8 infection (MOI 1; 24h) followed by immunoblotting with PCBP2 antibody. **(D)** RIP assay with Flag-PCBP2 in PR8 infected (MOI 1; 24h) HEK293T cells, enrichment of PCBP2 and circCPSF6 was analysed by immunoblotting with Flag antibody and RT-PCR, respectively. **(E)** Confocal microscopy image showing co-localization of circCPSF6 and PCBP2 in A549 cells. The cells were subjected immunostaining with PCBP2 antibody (green) followed by RNA-FISH with circCPSF6 specific Cy3 labelled probe (red), nuclei were stained with DAPI (blue). Scale bar, 10μm. The graph showing fluorescence intensity measured in ImageJ/Fiji, across cell. Data are presented as the mean ± SEM from triplicate samples of a single experiment and representative of two independent experiments. **** *p* < 0.0001 by two-tailed unpaired Student’s *t*-test. (Schematic is created with BioRender.com)

### CircCPSF6 modulates PCBP2–IPS-1 axis during IAV infection

Previous studies established PCBP2 as negative regulator of IPS-1 mediated antiviral signaling by promoting its proteasomal degradation^31^. Building on our finding that circCPSF6 interact with PCBP2, we further examine whether circCPSF6 modulate PCBP2 response during influenza virus infection. Supporting this, time course analysis indicated increase in protein expression of PCBP2, accompanied by increased transcript copy number quantified by dPCR in PR8 infected A549 cells **(Fig. 7A, B)**. Similar induction of PCBP2 transcript expression was observed by RT-PCR in PR8 infected human PBMCs and murine lung tissue **(Fig. 7C, D)**. Consistent with its reported function in other RNA viruses^31,32^, siRNA mediated knockdown of PCBP2 significantly reduced viral load in PR8 infected A549, human PBMCs, murine mpLFs and BMDCs **(Fig. S6A, Fig. 7E – G)**, underscoring proviral effect of PCBP2 during IAV infection. Mechanistically, PCBP2 silencing enhanced IPS-1 protein abundance in PR8 infected A549 cells **(Fig. S6B)**, consistent with PCBP2 mediated IPS-1 suppression.

**Fig 7.**
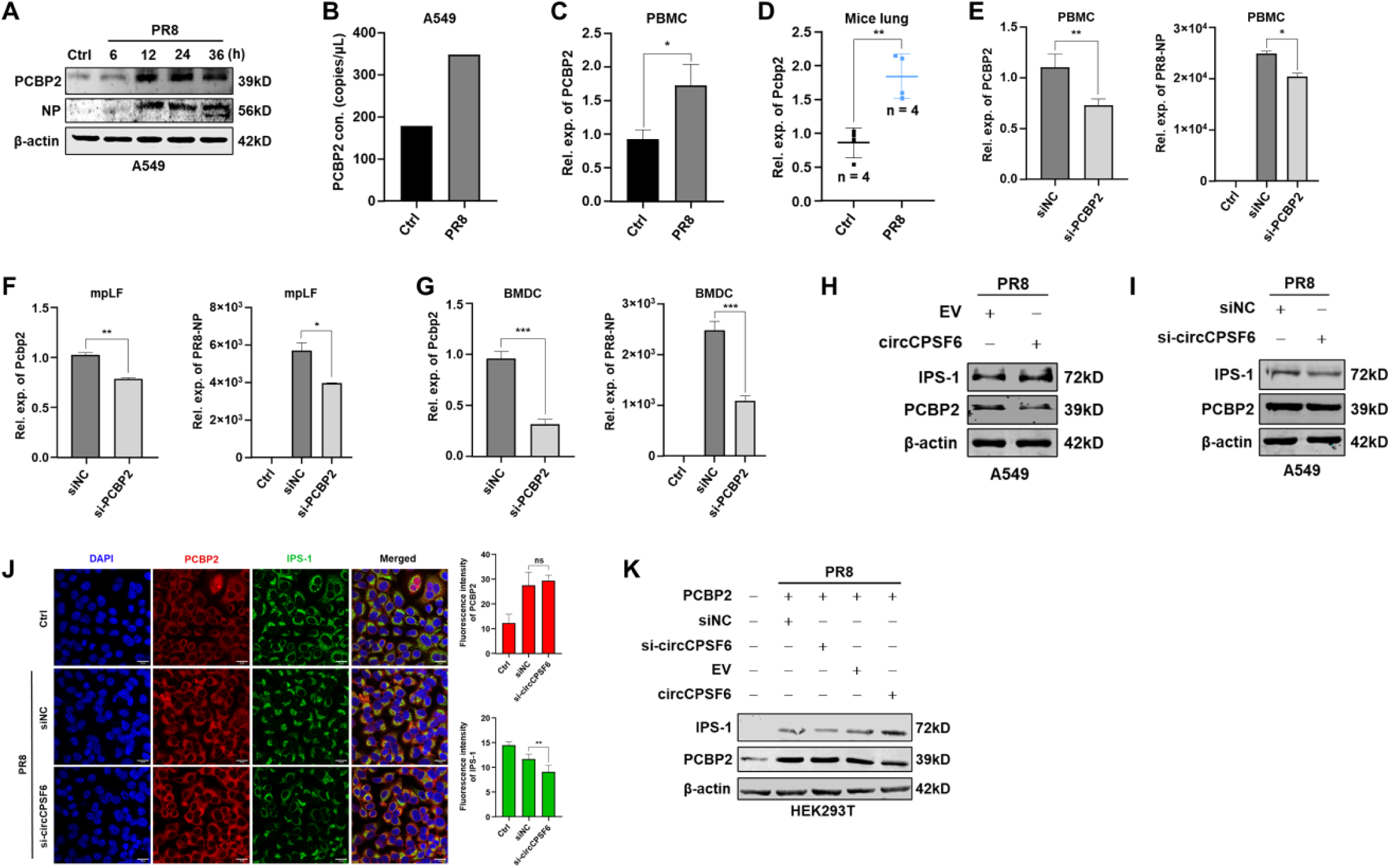
circCPSF6 regulates PCBP2 mediated IPS-1 degradation. **(A)** Immunoblotting of PCBP2 in PR8 infected (MOI 1) A549 cells at indicated time points. **(B)** Copy number of PCBP2 transcript determined by dPCR in PR8 infected (MOI 1; 24h) A549 cells. **(C)** Transcript level expression of PCBP2 determined by RT-PCR in PR8 infected (MOI 1; 24h) human PBMCs. **(D)** Transcript level expression of PCBP2 in lung tissue of mice infected with PR8 (PFU 100; 3 days), measured by RT-PCR. **(E – G)** Knockdown of PCBP2 and viral load measured by RT-PCR in PCBP2 knockdown **(E)** human PBMCs, **(F)** mice mpLFs and **(G)** BMDCs, followed by PR8 infection (MOI 1; 24h). **(H - I)** Immunoblot showing protein expression of IPS-1 and PCBP2 in **(H)** circCPSF6 overexpressing or **(I)** circCPSF6 knockdown A549 cells followed by PR8 infection (MOI 1; 24h). **(J)** Immunofluorescence with anti-PCBP2 (red) and anti-IPS-1 (green) in circCPSF6 knockdown, PR8 infected (MOI 1; 24h) A549 cells. Nuclei stain with DAPI (blue). Scale bar, 20μm. Bar graph representing fluorescence intensity of PCBP2 and IPS-1 protein. **(K)** Immunoblot showing protein level of IPS-1 and PCBP2 in HEK293T cells, co-transfected with PCBP2 and circCPSF6 knockdown or overexpression followed by PR8 infection (MOI 1; 24h). Data are presented as the mean ± SEM from triplicate samples of a single experiment and representative of three independent experiments. *** *p* < 0.001, ** *p* < 0.01, * *p* < 0.05, ns is non-significant by two-tailed unpaired Student’s *t*-test.

To define the role of circCPSF6 in this regulatory axis, we examined IPS-1 and PCBP2 protein expression in PR8 infected A549 cells. Ectopic expression of circCPSF6 markedly increased IPS-1 while reduced PCBP2 expression **(Fig. 7H)**. Conversely, circCPSF6 knockdown resulted decreased in IPS-1 protein level without affecting PCBP2 abundance, as shown by both immunoblotting and immunostaining **(Fig. 7I, J)**. Moreover, co-expression experiment in PR8 infected HEK293T cells, combined circCPSF6 silencing with PCBP2 overexpression resulted reduction of IPS-1 protein without affecting PCBP2 expression. Conversely, co-expression of circCPSF6 and PCBP2 resulted significant increase in IPS-1 and modest reduction of PCBP2 protein expression **(Fig. 7K)**.

Collectively these findings indicate that circCPSF6 engages PCBP2 to limit its abundance and modulate the PCBP2-IPS-1 axis by attenuating PCBP2 mediated degradation of IPS-1, thereby sustaining antiviral signaling during influenza virus infection.

## Discussion

Innate immunity provides the first line of host defence against virus infection by co-ordinated response of interferons and cytokines to restrict viral replication and prime adaptive immunity^33^. The precision of this system is regulated by complex networks of coding and non-coding elements of genome to equilibrate the antiviral response with immune homeostasis^34^. Our study identified non-coding circular RNA, circCPSF6, a previously unrecognized regulator of antiviral response, with a prominent role in context of RNA virus infection, particularly IAV. We demonstrated that circCPSF6 is highly conserved, predominantly localized in cytoplasm and significantly suppressed during viral infection both *in vitro* and *in vivo*. Functional analyses establish circCPSF6 as a critical barrier to viral replication, orchestrating robust induction of cytokines and antiviral genes.

Mechanistically, circCPSF6 act as a molecular sponge for miR-665 which is upregulated during IAV and other RNA virus infection. Our study shows, miR-665 function as a proviral factor by suppressing the expression of key antiviral mediators such as MyD88, STAT2 and IKKε of PRR-mediated antiviral innate immunity, thereby dampening host innate immune responses. CircCPSF6 act as ceRNA and relieve the repression of these antiviral mediators, hence, enhance cytokines and ISGs response. Beyond its miRNA sponging role, circCPSF6 interact with RBPs such as PCBP2 which is upregulated during IAV infection and act as a proviral mediator. PCBP2 is known to negatively regulate IPS-1 mediated antiviral signaling by enhancing its degradation. CircCPSF6 and PCBP2 interaction attenuate PCBP2-mediated IPS- 1 degradation by limiting the PCBP2 abundance, thereby sustaining IPS-1 levels and RLR-mediated antiviral signaling. This dual mechanism provides a new paradigm for highly sophisticated functional aspect of circRNA where circCPSF6 function as a central regulatory node by integrating both ceRNA network and RNA-protein interaction to fine-tune the innate immune response. This circCPSF6 mediated functional regulatory network represents a previously unrecognized layer of innate immune regulation that directly control the abundance of central adaptor proteins of PRRs. The integration of these regulatory networks enables a dynamic equilibrium of host innate immune system that ensure effective viral control with neutralizing excessive inflammation during virus infection.

Our findings highlight several broader concepts in host-virus interaction. First, it emphasizes the important role of circRNAs in innate immune response. The potential of circCPSF6 to function at both RNA-RNA and RNA-protein interface demonstrate the intricate balance of immune signaling at multiple levels by circRNAs. Such detailed regulation is crucial in virus infection as precise calibration of immune signaling prevent both viral replication and immune-mediated pathology. Second, it demonstrates the extend of viral exploitation of host non-coding RNAs. Several studies reported during IAV infection, some circRNAs such as circVAMP3^12^, circAIVR^10^, circMYO9A^11^, circCBP^13^ are upregulated to induce antiviral response, whereas, some circRNAs such as circ_0050463^35^, circ-GATAD2A^36^, circMerTK^37^ are upregulated to promote viral replication. Our study demonstrates consistent suppression of circCPSF6 and upregulation of miR-665 by variety of RNA viruses. Study showing differential expression profile of circRNAs in human lung epithelial cells during SARS-CoV-2 infection also report downregulation of circCPSF6 (hsa_circ_0000417) during infection, however its effect on SARS-CoV-2 virus is not explored^38^. This suggests a conserved viral strategy to disable the host antiviral factor. This highlights circCPSF6 as a crucial node in host defense and its significance in host-pathogen arm race within non-coding RNA networks.

Our study establishes circCPSF6 as a critical modulator of innate antiviral immunity, however, there are certain limitations remain to be explored. The mechanistic analysis was primarily done *in vitro* cell culture model, however, *in vivo* functional study of circCPSF6 in knockout or overexpression animal models are required to dissect the physiological relevance during virus infection. Study in esophageal squamous cell carcinoma shows that circCPSF6 can interact with its parent gene CPSF6 which is an RNA binding protein^39^. During virus infection, whether this interaction occur and contribute to circCPSF6 function or regulation remains an open question. Moreover, the mechanistic analysis of this study based on IAV infection. However, this study can be extended in other RNA viruses to determine the broader spectrum of circCPSF6 mediated antiviral response.

From a translational perspective, our findings open up a new avenue for RNA-based therapeutic possibilities. The inherent stability of circRNA makes them highly attractive for both diagnostic biomarkers and potential therapeutic agents^40^. The mechanistic findings of our study indicate, restoring circCPSF6 level, either through synthetic circRNA delivery or by inhibiting viral mechanism that mediate its suppression could represent a potential pan-antiviral strategy. Such approach could complement the existing antiviral therapies by reinforcing intrinsic host defence network making it more resilient to viral escape strategies, effective across multiple viral species and potentially reduce the risk of viral resistance. This study deepens our understanding of the complex network of innate immune regulation and provide an intriguing concept for development of new immunomodulatory therapies.

## Acknowledgements

We thank R. Fouchier for providing the A/PR8/H1N1 reverse genetics system. We acknowledge the Central Instrumentation Facility at IISER Bhopal for technical support. We are grateful to Dr. Anamika Mishra and Dr. Ashwin A. Raut (ICAR-NIHSAD) for their assistance with virus propagation. We thank Kanishk Sharma and Pratik Katekar for technical support. This work was partially supported by the Indian Council of Medical Research (ICMR; Grant No. ICMR/BIO/2023-2024/98) and by institute funding from IISER Bhopal. The funders had no role in study design, data collection and interpretation, or the decision to submit the work for publication. A.M. thank DBT for awarding SRF. R.C thank UGC for awarding SRF.

## Contributions

**Aparna Meher** – Conceptualization, Methodology, Validation, Formal Analysis, Investigation, Data curation, Animal experiments, Writing – Original Draft, Writing- Review & Editing, Visualization

**Riya Chaudhary** – Methodology, Validation, Formal Analysis, Investigation, Animal experiments, Writing- Review & Editing, Visualization

**Himanshu Kumar** – Conceptualization, Methodology, Formal Analysis, Data curation, Writing – Original Draft, Writing- Review & Editing, Project administration, Funding Acquisition, Visualization, Supervision

## Ethics declaration

All animal experiments were performed in compliance with institutional guidelines and approved by the Institutional Animal Ethics Committee (IAEC) of IISER Bhopal (Approval No. 2025-IISERB-7.9-IAEC). All experiments involving influenza virus and Newcastle disease virus were conducted under biosafety guidelines with approval from the Institutional Biosafety Committee (IBSC) of IISER Bhopal (Approval No. IBSC/IISERB/2025/Meeting- 1/05).

---

**Supplementary Fig 1.**
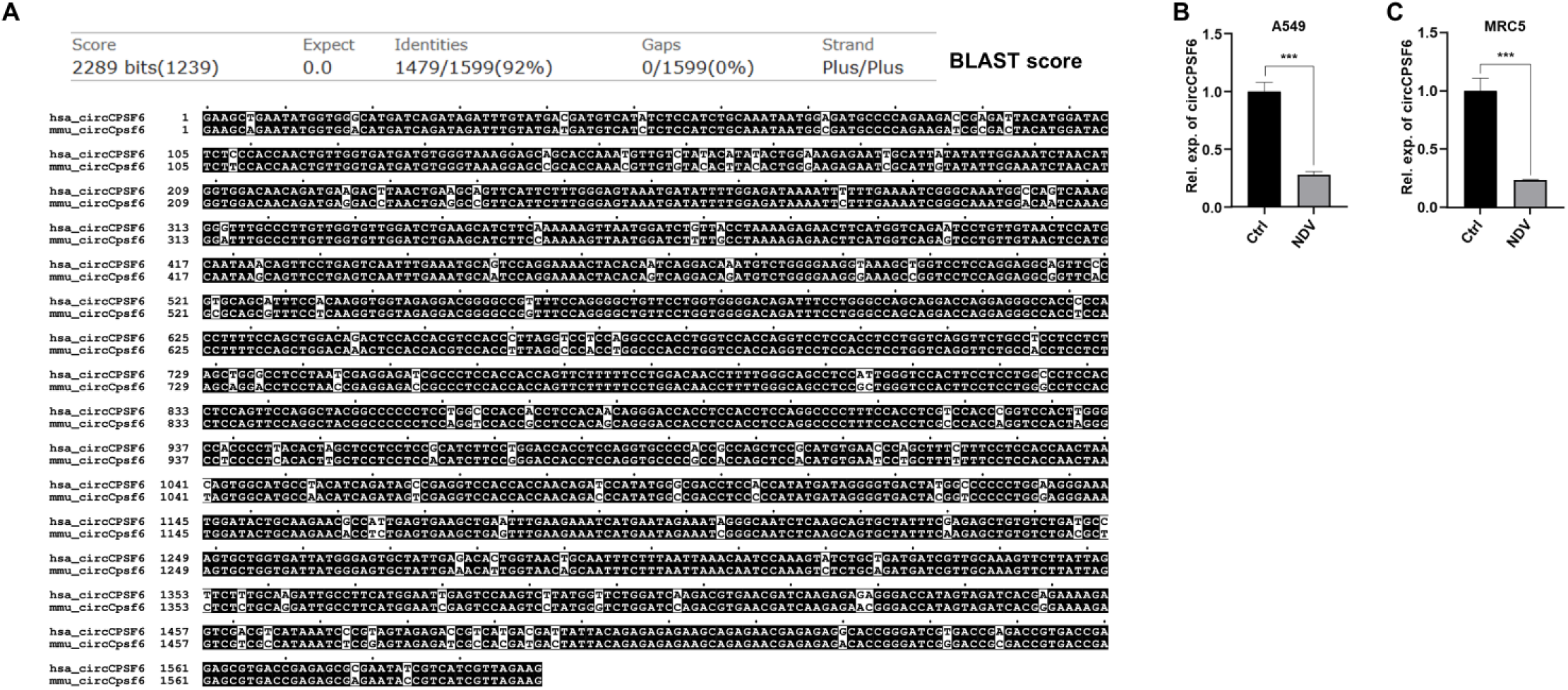
CircCPSF6 conservation and expression during RNA virus infection. **(A)** Sequence alignment of hsa_circCPSF6 and mmu_circCpsf6 was performed using basic local alignment search tool (BLAST). **(B - C)** CircCPSF6 expression level determined by RT-PCR in NDV infected (MOI 1; 24h) **(B)** A549 cells and **(C)** MRC5 cells.

**Supplementary Fig 2.**
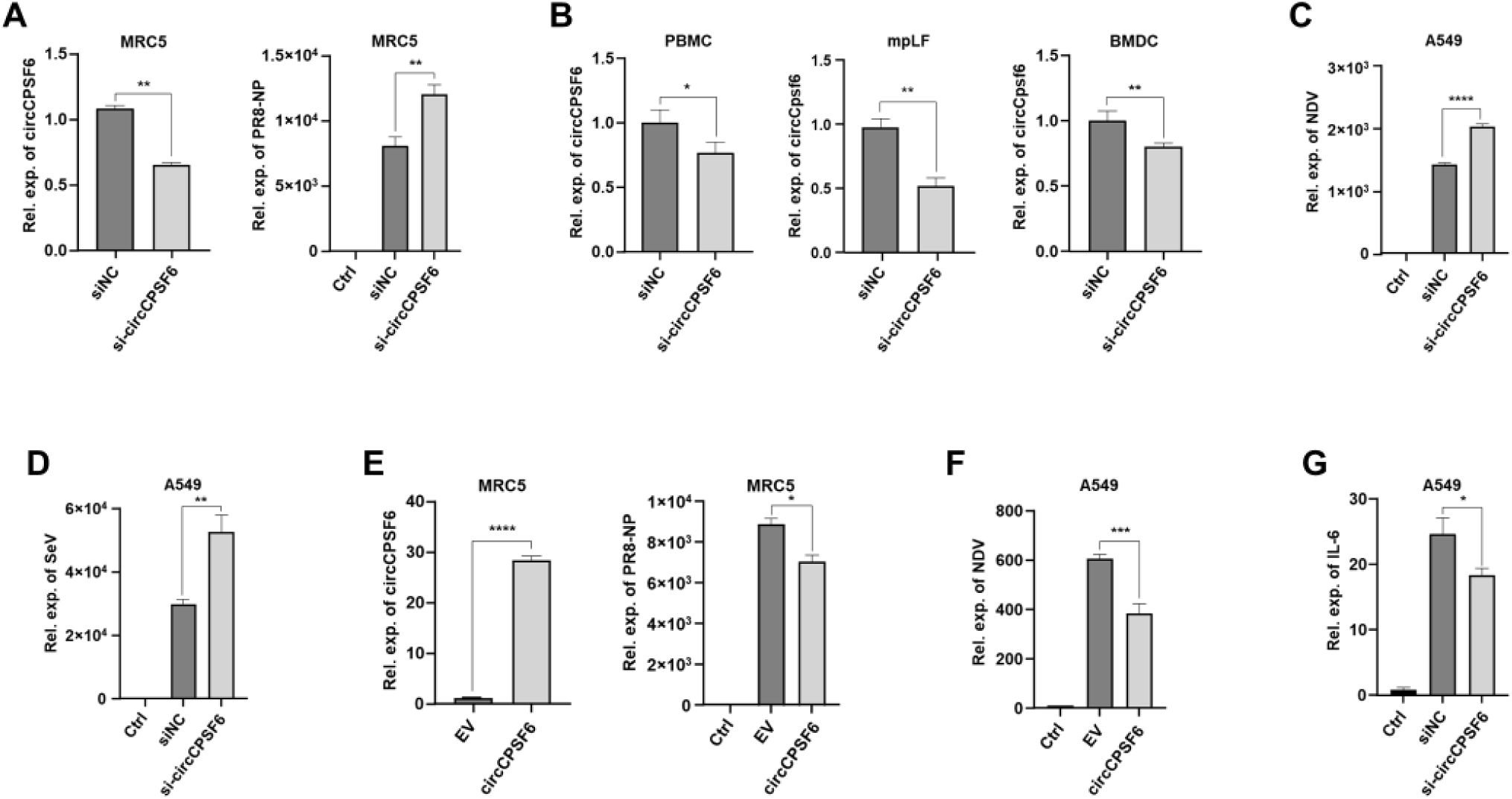
CircCPSF6 inhibit RNA virus replication. **(A)** CircCPSF6 knockdown and viral load was measured by RT-PCR in PR8 infected (MOI 1; 24h) MRC5 cells. **(B)** Knockdown of circCPSF6 measured by RT-PCR in PR8 infected (MOI 1; 24h) human PBMCs, mice mpLFs and BMDCs. **(C – D)** Viral load measured by RT-PCR in circCPSF6 knockdown A549 cells followed by **(C)** NDV infection (MOI 1; 24h) and **(D)** SeV infection (MOI 2; 24h). **(E)** Overexpression level of circCPSF6 and viral load measured by RT-PCR in PR8 infected (MOI 1; 24h) MRC5 cells. **(F)** Viral load measured by RT-PCR in circCPSF6 overexpressing, NDV infected (MOI 1; 24h) A549 cells. **(G)** IL-6 transcript expression measured by RT-PCR in circCPSF6 knockdown, PR8 infected A549 cells (MOI 1; 24h).

**Supplementary Fig 3.**
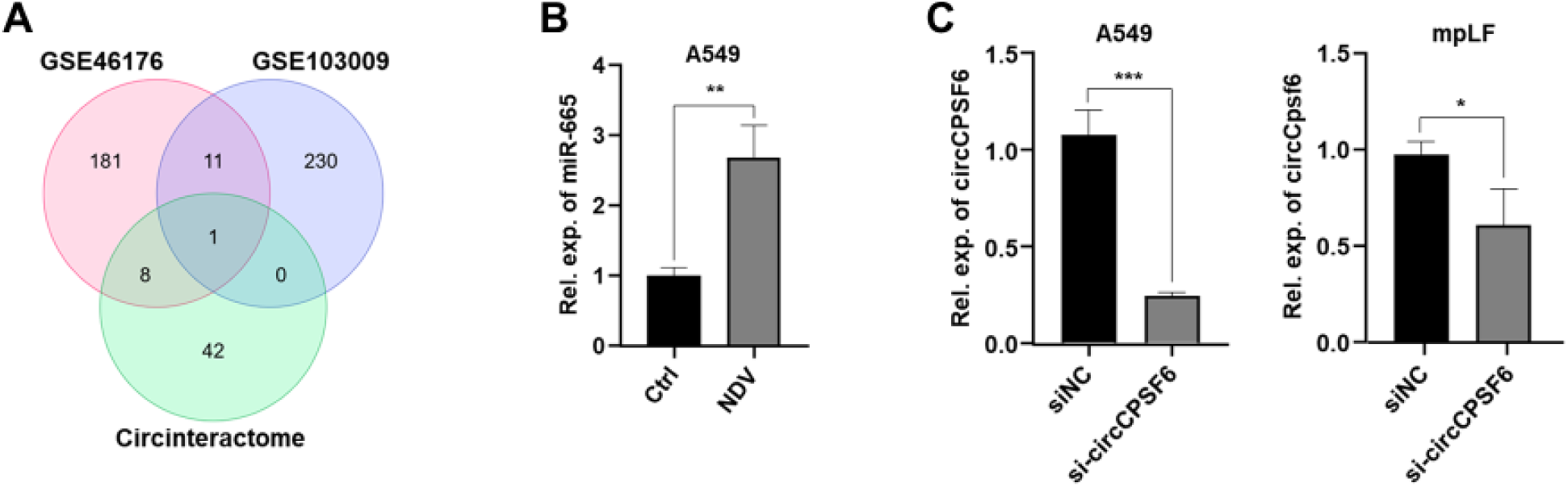
Screening and expression of miR-665 during RNA virus infection. **(A)** Venn diagram showing common miRNA found by overlapping GEO dataset result and miRNAs identified in Circinteractome. **(B)** Expression level of miR-665 measured by RT-PCR in NDV infected (MOI 1; 24h) A549 cells. **(C)** Knockdown of circCPSF6 measured by RT-PCR in PR8 infected (MOI 1; 24h) A549 cells and mpLFs.

**Supplementary Fig 4.**
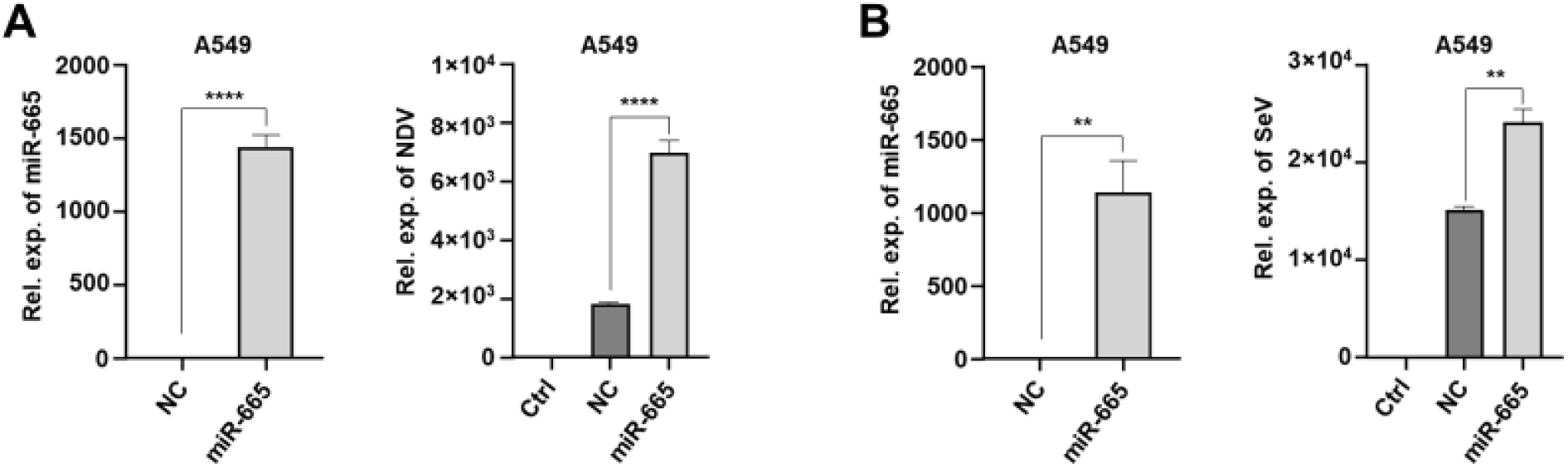
miR-665 enhance RNA virus replication. **(A)** Overexpression of miR-665 mimic and viral load determined by RT-PCR in NDV infected (MOI 1; 24h) A549 cells. **(B)** Overexpression of miR-665 mimic and viral load determined by RT-PCR in SeV infected (MOI 2; 24h) A549 cells.

**Supplementary Fig 5.**
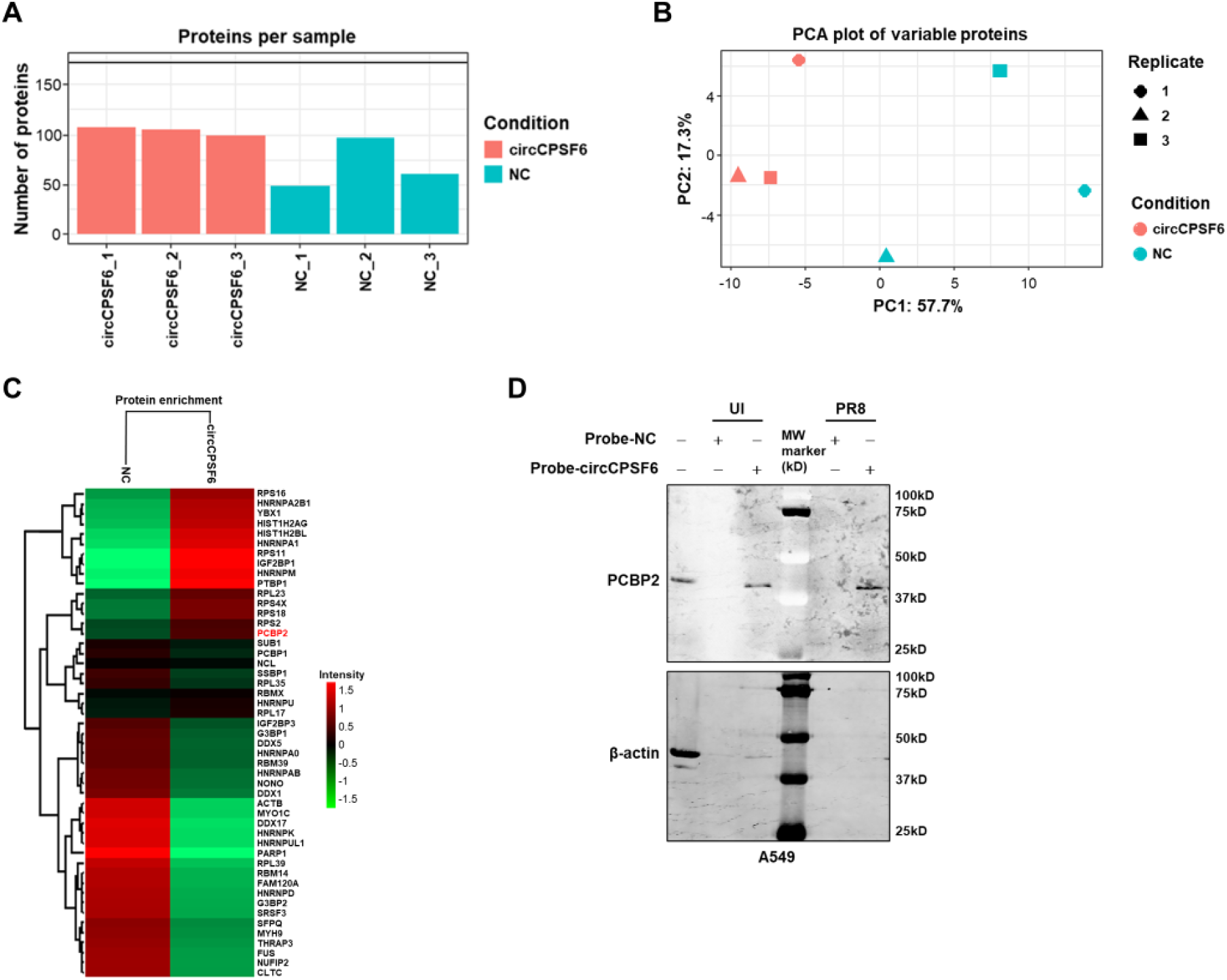
Identification of circCPSF6 interacting proteins. **(A)** Graph showing number of proteins identified in circCPSF6 probe fraction by mass spectrometry. **(B)** Principal component analysis (PCA) of normalized protein intensities identified in circCPSF6 probe and NC probe fraction. Each point indicates one individual sample replicate from distinct cluster based on variance (PC1, 57.7%; PC2, 17.3%). **(C)** Heatmap showing mean normalized protein enrichment in circCPSF6 probe and NC probe pulldown fraction. Colour scale represents normalized enrichment based on Z-score (red, high; green, low). **(D)** Raw image of RNA pulldown assay with biotin labelled probe specific to circCPSF6 BSJ in A549 cells with or without PR8 infection (MOI 1; 24h) followed by immunoblotting with PCBP2 antibody.

**Supplementary Fig 6.**
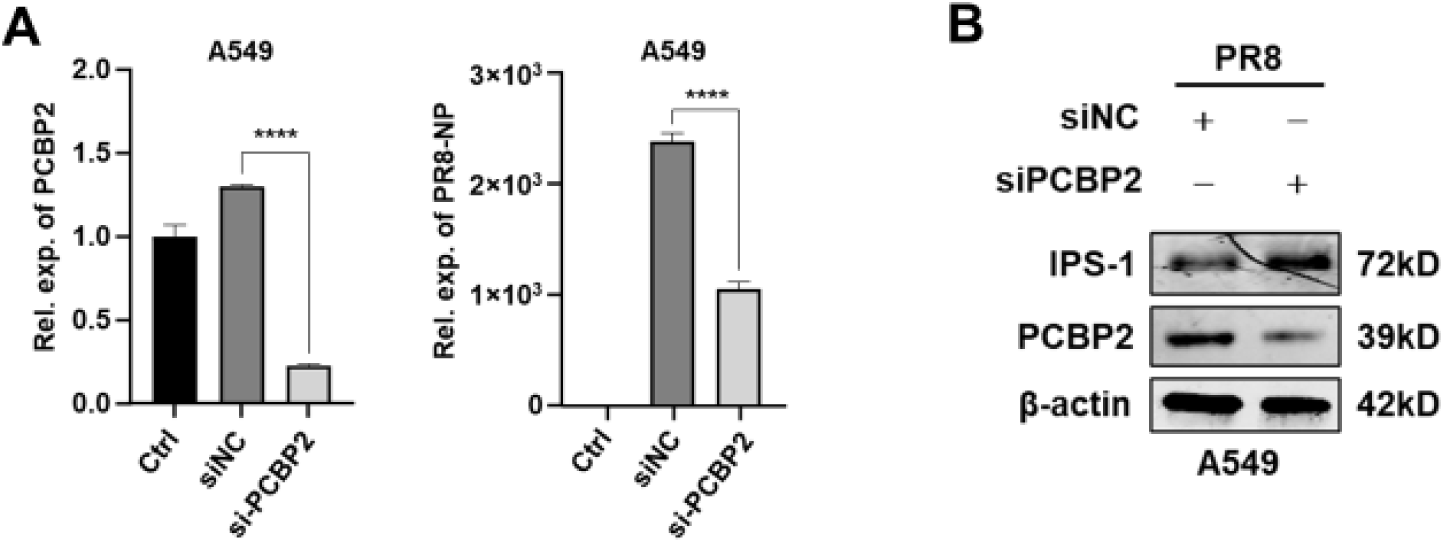
CircCPSF6 modulate PCBP2-IPS-1 axis. **(A)** Knockdown of PCBP2 and viral load determined by RT-PCR in PCBP2 knockdown, PR8 infected (MOI 1; 24h) A549 cells. **(B)** immunoblot showing PCBP2 and IPS-1 protein expression level in PCBP2 knockdown, PR8 infected (MOI 1; 24h) A549 cells.

**Supplementary Table 1.** Predicted binding sites of miR-665 on hsa_circCPSF6.

Predicted binding sites of miR-665 on hsa\_circCPSF6
| miRNA | No. of sites | Position on circCPSF6 | Score | Energy (kCal/Mol) |
| --- | --- | --- | --- | --- |
| miR-665 | 1 | 689 - 708 | 166 | -30.19 |
|  | 2 | 806 - 825 | 158 | -26.82 |
|  | 3 | 847 - 864 | 154 | -24.36 |
|  | 4 | 956 - 975 | 143 | -19.93 |

**Supplementary Table 2.** Predicted binding sites of miR-665 on mmu_circCpsf6.

| **miRNA** | **No. of sites** | **Position on circCpsf6** | **Score** | **Energy (kCal/Mol)** |
| --- | --- | --- | --- | --- |
| miR-665 | 1 | 689 - 708 | 166 | -30.19 |
|  | 2 | 806 - 825 | 158 | -26.74 |

**Supplementary Table 3.**
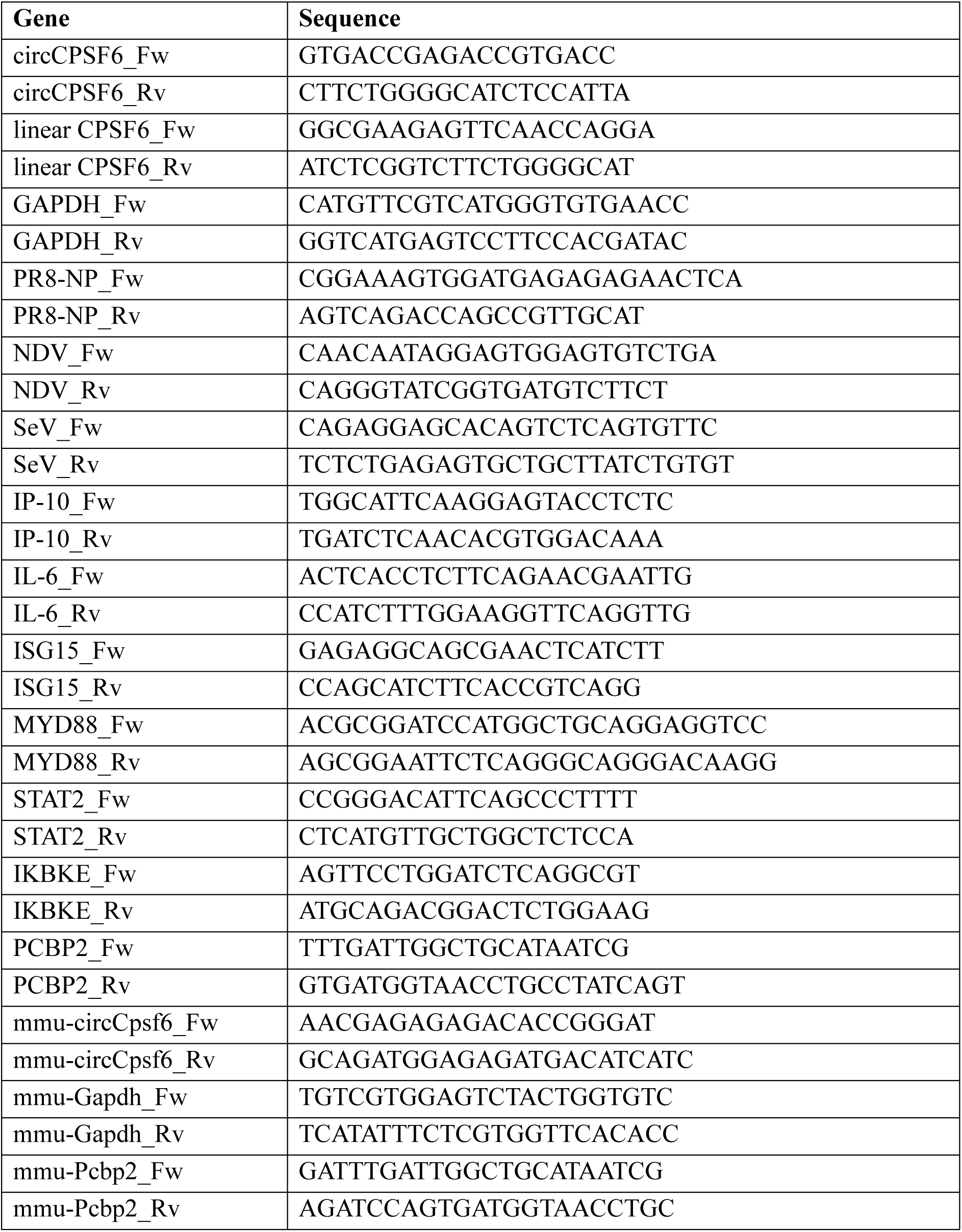
Primer list for RT-PCR.

